# Sensor sensibility: Divergent measurements of dopaminergic signaling to acute morphine administration via fiber photometry

**DOI:** 10.64898/2026.06.19.733408

**Authors:** Rachel M. Donka, Maxine K. Loh, Mitchell F. Roitman, Jamie D. Roitman

**Author notes:** Corresponding author: Jamie D. Roitman 1007 W Harrison St. Chicago IL 60607.

## Abstract

Activity of the mesolimbic dopamine system has long been implicated in encoding primary rewards and contributing to the addictive properties of drugs of abuse. Dopamine neurons in the ventral tegmental area (VTA_DA_) of the midbrain typically show patterns of spontaneous burst activity that align with the onset of salient events or rewarding stimuli, resulting in phasic dopamine release in the nucleus accumbens (NAc). Fiber photometry is increasingly being used as an accessible technique to quantify neural activity with high temporal resolution at sensors offering signal specificity in stable recordings over extended periods of time. It has been well established by multiple techniques that opioids increase mesolimbic dopamine activity, likely through disinhibition of VTA_DA_ neurons. Here we used fiber photometry to compare sub-second transient events from VTA_DA_ neurons with GCaMP6f and dopamine release in the lateral shell of the NAc with dLight1.3b and GRABDA2h in response to morphine treatment. In weekly sessions, one dose of morphine was administered in escalating order (2.5, 5,7.5, and 10 mg/kg, intraperitoneal). Consistent with prior literature, both GCaMP6f in VTA_DA_ neurons and dLight1.3b in NAc showed patterns of increased signal following morphine treatment. In contrast, morphine suppressed transient activity at GRABDA2h sensors. Further analyses of whole signal streams from each sensor showed a generalized increase, but reduction in variability of the GRABDA2h signal, consistent with the interpretation of sensor saturation. Such results emphasize the importance of the inclusion of appropriate controls to contextualize the interpretation of biosensor responses, particularly in response to pharmacological treatment.

**HIGHLIGHTS:**

- Morphine elicited increased signaling in VTA_DA_ GCaMP6f and NAc dLight1.3b, consistent with prior literature
- Morphine suppressed NAc GRABDA2h signaling of transient events, suggesting saturation of GRABDA2h sensor
- Sensor validation with pharmacological challenges is critical for interpretation of data

## 1. INTRODUCTION

Primary rewards briefly evoke high frequency activity of dopamine (DA) neurons (Schultz, 1997) and high concentration of DA release in the nucleus accumbens (NAc; (Roitman et al., 2004)). These phasic events, termed transients, also develop to reward predictive cues (Amo et al., 2022; Burke et al., 2026) and are thought to be critical for the learning of reward-related relationships. They also occur ‘spontaneously’ - that is, without any clear, driving stimulus (Owesson-White et al., 2012). A common property of drugs of abuse is to increase the frequency and amplitude of dopamine transients (Cheer et al., 2007; Covey et al., 2014), although do so via different mechanisms (Lüscher & Ungless, 2006). It is well established that the opioid morphine increases dopamine transients in the NAc (Fox et al., 2016; Vander Weele et al., 2014). Indeed, the reinforcing effects of opioids are substantially mediated at mu opioid receptors (MOR), which are concentrated on GABAergic interneurons in the ventral tegmental area (VTA) that provide inhibitory tone on VTA DA (VTA_DA_) neurons (Garzón & Pickel, 2001; Le Merrer et al., 2009; Negus et al., 1993). Activation of MORs reduces inhibition of DA cell bodies (Johnson & North, 1992) thereby releasing suppression of VTA_DA_ activity (Bourdy & Barrot, 2012; Jalabert et al., 2011).

Seminal work describing the effects of morphine on phasic dopamine signaling was established with electrophysiological recordings from presumed DA cell bodies or fast-scan cyclic voltammetry in the NAc. However, corroborative studies are needed as electrophysiological signatures of DA neurons are not reliable (Margolis et al., 2010), and much heterogeneity among dopamine neurons has been elucidated (Blaess & Krabbe, 2023; Phillips et al., 2022). With respect to measuring release, fast-scan cyclic voltammetry lacks ideal analyte specificity and is subject to pH shifts and movement artifacts (Rodeberg et al., 2017), creating a need for DA recording methods with greater specificity and motion control considerations.

The development of fiber photometry for *in vivo* neural recordings has rapidly been adopted due to its fast temporal sampling, outstanding specificity, high yield, and ability to account for signal artifacts. The introduction of fiber photometry (Gunaydin et al., 2014) paired with transgenic animals (Witten et al., 2011) to selectively express genetically encoded calcium (Ca^2+^) sensors enables recording from DA cell bodies using changes in fluorescence as a proxy for cell body activity (Simpson et al., 2024). Biosensors can also selectively measure DA release (Chen et al., 2013; Helassa et al., 2016) via modified DA receptors including dLight (modified D_1_ receptor (Condon et al., 2021; Patriarchi et al., 2018)) and GRABDA (modified D_2_ receptor (Sun et al., 2020)). Using fiber photometry with DA specific biosensors allows for analysis of spontaneous events, gradual shifts over time, and precise responses to stimuli (Labouesse et al., 2020). While multiple studies have used fiber photometry to assay DA activity in response to naturalistic stimuli (Hsu et al., 2020) and pharmacological treatments including opioid administration (Çimen & Kutlu, 2025; Culver et al., 2026; Higginbotham et al., 2025; Lefevre et al., 2020), validation of performance across sensors is limited. Here we establish the acute effects of escalating doses of morphine on DA activity, recording VTA_DA_ cell body activity via GCaMP6f as well as downstream DA release in NAc via dLight1.3b and GRABDA2h. We sought to identify the temporal dynamics of changes in DA signaling in response to differing doses of morphine, validate the alignment in experimental outcomes between cell body activity and DA release, and assay fidelity across multiple biosensors.

## 2. MATERIALS AND METHODS

### 2.1. Subjects

Male (n=13) and female (n=13) Long Evans rats (>250 g) were bred from heterozygous females expressing Cre recombinase under the control of the tyrosine hydroxylase promoter (TH:Cre+; Rat Research Resource Center, RRRC No. 659; Witten et al., 2011) and male wildtype Long Evans Rats (Charles River Laboratories). Upon weaning at postnatal day (PD) 22, offspring were genotyped for Cre expression (Transnetyx, Inc.) and pair- or group-housed until PD60. Rats were then pair-housed in a temperature- and humidity-controlled vivarium on a 12:12 hour light:dark cycle (lights on at 0700 hour) and provided with *ad libitum* food and water for the duration of all experiments. Experimental testing was conducted in a separate room during the light phase of the cycle. All studies were conducted in accordance with the National Institutes for Health Guide for the Care and Use of Laboratory Animals and approved by the Animal Care Committee (ACC) at the University of Illinois Chicago.

### 2.2. Viruses

*In vivo* fiber photometry experiments used adeno-associated viruses (AAVs) packaged with fluorescent protein sensors. To detect intracellular calcium in TH+ cell bodies, we used AAV1.hSyn.Flex.GCaMP6f.WPRE.SV40 (n=8; 5×10^12^ GC/mL, Addgene; (Zhang et al., 2023)). To detect dopamine, we used AAV9.hSyn.GRABDA2h (n=9; 1×10^13^ GC/ml, Addgene; (Sun et al., 2020)) and AAV9.syn.dLight1.3b (n=9; 1×10¹³ vg/mL, Addgene; (Patriarchi et al., 2018)).

### 2.3. Surgical Procedures

Rats were anesthetized with isoflurane (4% induction, 2% maintenance) and injected with systemic (1mg/kg meloxicam, subcutaneous (s.c.)) and local analgesic (2mg/kg bupivacaine, s.c.). Depth of anesthesia was assessed via toe pinch and eye blink reflex periodically through the procedure. The incision site was cleared of hair and sterilized with alternating betadine and 70% ethanol washes, and the animal was secured in a stereotaxic frame (Kopf Instruments). Stereotaxic measurements were made relative to Bregma (Paxinos & Watson 1998). To record VTA_DA_ activity, AAV1.hSyn.Flex.GCaMP6f.WPRE.SV40 (1 µl; 0.1 µl/minute, followed by 5-minute post-injection period for diffusion) was injected via a Hamilton syringe (10 µL Gastight Syringe, #7653-01) and targeted to the VTA (AP -5.4; ML +/- 0.6; DV -8.15 mm; Fig. 1A). An optic fiber (flat 400-μm core, 0.48 numerical aperture [NA], Doric Lenses Inc.) was implanted in the VTA immediately dorsal to the injection site (AP -5.4; ML +/- 0.6; DV -8.00 mm). To measure dopamine release, 1 µl of AAV9.hSyn.GRABDA2h or AAV9.syn.dLight1.3b was injected targeting the NAc lateral shell (AP: +1.5; ML: +/-2.5; DV: –8.0 mm) and an optic fiber was implanted dorsal to the injection site (DV: -7.9 mm; Fig. 1B,C). Optic implants were secured to the skull by three stainless steel screws covered by Metabond (Parkell, Inc) and dental acrylic (Patterson Dental Company). Meloxicam (1 mg/kg, s.c.) and physiological saline (10 ml/kg, s.c.) were administered for two days following surgery. Postoperative monitoring of the incision site and subject body weight was conducted for a minimum of 10 days. To allow for recovery and construct expression, experiments began at minimum 28 days after surgery.

**Figure 1.**
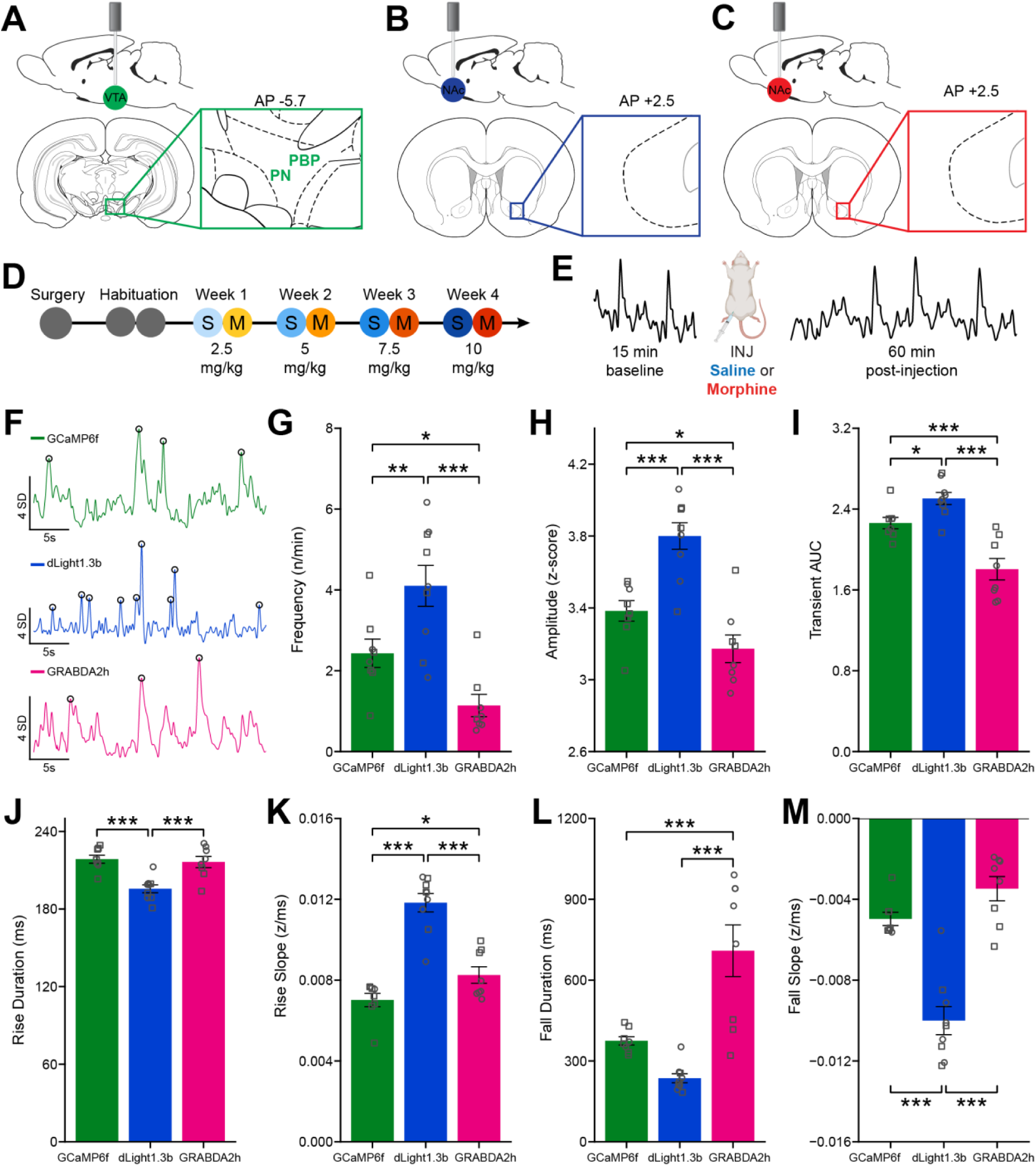
Experimental methods and baseline sensor spontaneous transient comparisons. Sensor and fiber optic implant targeting for **A)** Cre-dependent GCaMP6f in the VTA in TH-Cre Long Evans rats to record dopamine cell body activity, **B)** dLight1.3b in the nucleus accumbens lateral shell (NAcLS), and **C)** GRABDA2h in the NAcLS. **D)** Experimental timeline. Recordings began a minimum of 4 weeks post surgery. Subjects were habituated to operant chambers for two days prior to the dose-response paradigm. Morphine doses were administered once weekly in escalating order from 2.5 to 10 mg/kg, with a saline control session performed on the day prior to morphine treatment each week. **E)** Session design. Each session consisted of a 15-minute baseline period (first 3 minutes excluded from analysis to account for signal stabilization and photobleaching), i.p. injection of saline or morphine, and a 60-minute post-injection recording period. **F)** Example traces with identified spontaneous transient events by sensor. Circles mark identified transients. The first saline session was used for baseline transient parameter quantification and comparison between sensors. Mean transient parameter comparisons between sensors: **G)** frequency, **H)** amplitude, **I)** area under the curve, **J)** rise duration from half-height to peak, **K)** rise slope from half-height to peak, **L)** fall duration from peak to half-height, and **M)** fall slope from peak to half-height. * <0.05; **<0.01; ***<0.001

### 2.4. Experimental Paradigm

Over 4 weeks, animals completed two recording sessions on consecutive days each week to test the acute effects of morphine administration on fiber photometry signaling. All subjects were tested with four acute doses of morphine sulfate (10 mg/ml; intraperitoneal (i.p.)): 2.5, 5.0, 7.5, and 10.0 mg/kg. For each dose, we performed a control session (0.9% sterile saline; volumetrically matched to morphine sulfate dose) followed by a morphine treatment session on the next day (Fig. 1D). Doses were administered in escalating order to mitigate the possibility of hold-over and tolerance effects (Zhang et al., 2008). Within each session, a 15-minute pre-injection baseline was recorded, followed by time-stamped injection of saline control or morphine sulfate, followed by an additional 60 minutes of post-injection recording (Fig. 1E).

### 2.5. Equipment

Fiber photometry recordings were conducted in accordance with prior protocols (Donka et al., 2025). LEDs (Tucker Davis Technologies) administered 465 nm (Ca^2+^-dependent, dLight1.3b dopamine-dependent, or GRABDA2h dopamine-dependent) excitation and 405 nm control wavelengths. Output power was set for each subject prior to the first recording session between 30 and 50 µW output at each wavelength. Power levels were held constant for the duration of all experiments. Intensity of the 465 nm and 405 nm light was sinusoidally modulated at 211 and 531 Hz respectively. Light was coupled to a filter cube (FMC4, Doric Lenses), converged into a pigtailed optical fiber rotary joint (FRJ_1x1_PT-G2, Doric Lenses), and connected to the fiber optic implant of the animal via an optical fiber patch cord (MFP_400/430/LWMJ-0.57_.47m_FCM-MF2.5 (F)_LAF, Doric Lenses). Fluorescence was collected via the same patch cord and focused into a photoreceiver (Visible Femtowatt Photoreceiver Model 2151, Newport). A lock-in amplifier and data acquisition system (RZ5P or RZ10X; Tucker Davis Technologies) was used to demodulate the fluorescence due to 465 nm and 405 nm excitation, and data were stored using the data acquisition software Synapse (Tucker Davis Technologies).

Experimental testing was conducted in standard operant conditioning chambers (ENV-009A-CT; Med Associates Inc.) outfitted with a houselight and fan. Operant chambers were enclosed by sound attenuating boxes (ENV-016MD; Med Associates Inc.). Injections during recording sessions were timestamped at the initiation and completion of injection via manually generated TTLs (DIG-726; Med Associates Inc.) and sent to the fiber photometry data acquisition system and stored as TTLs by the recording software Synapse (Tucker Davis Technologies).

### 2.6. Immunohistochemistry

Following completion of all experiments, rats were deeply anesthetized with isoflurane and transcardially perfused with 0.9% NaCl followed by a 10% buffered formalin solution (HT501320, Sigma-Aldrich). Brains were removed and stored in formalin with 20% sucrose. Brains were sectioned at 40 µm on a freezing stage microtome (SM2010R, Leica Biosystems). Sections were collected and processed to label GFP as an indicator of sensor expression and/or TH (VTA GCaMP6f recordings) via immunohistochemistry. Free-floating tissues were washed in 1x potassium-phosphate buffered solution (KPBS) six times for 10 minutes each, permeabilized in 0.3% Triton X-100 in KPBS for 30 minutes, and blocked in 2% normal donkey serum for 30 minutes. Sections were incubated in primary chicken anti-GFP (AB13970, Abcam) and/or rabbit anti-TH (AB152, Sigma-Aldrich) antibodies overnight (∼18 hr) at 4° C. Sections were washed six times for 10 minutes each and incubated in overnight in secondary antibody (AF488 conjugated donkey anti-chicken and/or Cy3 conjugated donkey anti-rabbit; Jackson Immunoresearch). Sections were washed, mounted onto glass slides, air dried, and cover slipped with Fluoroshield with DAPI (F6057, Sigma-Aldrich). To verify viral expression and fiber implant localization, tissue sections were visualized and photographed with bright-field microscopy (Olympus BX43 Fluorescence Research Microscope). Data from rats with GFP expression and fiber placements within the borders of the VTA or NAc were included in analyses.

### 2.7. Data Analyses

#### 2.7.1. Fiber Photometry Signal Processing

Fiber photometry signal processing was conducted in MATLAB (R2025a) using the open-source PASTa protocol and toolbox v1.1.0 (Donka et al., 2025). Briefly, raw 465 nm and 405 nm streams and injection start and end TTLs were extracted into MATLAB. Prior to signal processing, the first three minutes of the recording session, during which most photobleaching occurs, was trimmed and event TTLs were adjusted to maintain temporal alignment. Raw 405 nm streams were scaled to the 465 nm by a constant scaling factor determined based on the frequency domain power spectrum. Background scaling frequency minimums (GCaMP6f: 10 Hz; dLight1.3b: 10 Hz; GRABDA2h: 8 Hz) and maximums (GCaMP6f: 100 Hz; dLight1.3b: 100 Hz; GRABDA2h: 50 Hz) were set optimally for each sensor to the range that minimized over and under scaling of the 405 nm stream relative to the 465 nm stream. For each session, raw 405 nm background stream was multiplied by the scaling factor and centered around the mean of the 465 nm stream. To account for photobleaching and motion artifact, the 465 nm stream was converted to ΔF/F via subtraction of the scaled 405 nm stream from the 465 nm stream, followed by division by the scaled 405 nm stream.

#### 2.7.2 Transient Event Detection

Prior to transient event detection, the subtracted signal was filtered with a third order bandpass Butterworth filter (highpass cutoff frequency: 0.0051 Hz; lowpass cutoff frequency: 2.286 Hz). Whole session subtracted and filtered signals were Z-scored to the mean and standard deviation of the 12-minute pre-injection baseline within each session. Individual transient events were detected as described in Donka et al, 2025 (Fig. 1F). Briefly, event inclusion threshold was set at 2.6 standard deviation (SD) based on the pre-injection baseline period. To detect individual events, each peak in the data stream was localized and the maximum value (peak max) was compared to a pre-event baseline value, determined as the mean value in the time window of 1000 to 100 ms pre-peak max. Events that exceeded threshold were included as transient events for analysis.

Once detected, the frequency, amplitude, and area under the curve (AUC) of individual transient events were quantified in each three-minute time bin of the post-injection period. Frequency was measured as the number of transient events per minute in each bin. Amplitude was calculated as the difference between the peak max and baseline value of the event. AUC was calculated as the trapezoidal AUC between the baseline and observed z-score from 100 ms pre-peak max to 1500 ms post-peak max. Quantification of AUC was included to examine gross changes in transient shape that might not be captured in measurement of frequency or amplitude. Additional quantification metrics are included in the supplemental data files.

#### 2.7.3. Whole Session Quantification

In addition to analyses of transient events, we quantified the mean and variation of the whole subtracted stream of data from 12 minutes pre-injection through 60 minutes post-injection. To facilitate comparisons of shifts in signal mean across the session between sensor groups, subtracted signal streams were normalized (Z-score) to the mean and standard deviation of the pre-injection period for each session. Overall mean and mean absolute deviation (MAD, mean absolute value of the distance between each data point and the mean) of each stream were calculated in three-minute time bins across the session (4 bins pre-injection, 15 bins post-injection). Post-injection time bin MAD values were scaled to pre-injection baseline MAD to facilitate comparisons between subjects and sensor groups:

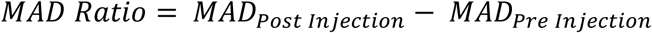

#### 2.7.4. Statistical Analyses

All statistical analyses were conducted in R (R Core Team, 2023) and all analysis scripts are available at https://github.com/rdonka/SensorSensibility.

##### Individual session analyses

To quantify the effect of morphine treatment on the frequency of transient events, frequency was calculated as transients per minute within each three-minute time bin across the session. For amplitude and AUC, transients were divided into time bins and models were fit to all individual events. Generalized linear mixed effects models were fit with glmmTMB (Brooks et al., 2017) separately for each sensor.

We first fit a Gaussian mixed-effects model with fixed effects and interactions between treatment (saline or morphine), dose (2.5, 5.0, 7.5, or 10 mg/kg), and time bin (4 pre-injection and 20 post-injection) with random intercepts by subject. Because repeated time bin measurements within subjects may exhibit temporal autocorrelation, we fit a second model with a first order autoregressive (AR(1)) correlation structure applied to time bins within each subject by injection type. This specification accounts for correlation among nearby time points, providing more accurate estimates when residuals are temporally dependent. Model fit was compared using Akaike’s Information Criterion (AIC), with lower AIC values indicating better fit while penalizing model complexity. Results for model fit are specified in the supplementary tables. Statistics reported are based on the AR(1) model when it provided a lower AIC than the base model, otherwise results from the base model are reported. Type II Wald Χ^2^ tests (Fox & Weisberg, 2019) were used to evaluate fixed-effect terms.

All post-hoc tests were conducted using emmeans (Lenth et al., 2024). For each model, we tested the joint significance of levels of injection type and dose to assess significant differences among levels of each factored variable. Significant joint tests by injection type or dose were followed with pairwise simple-effects comparisons. Results from these analyses are shown in Figures 2, 3, and 4, panels C, J, and Q.

**Figure 2.**
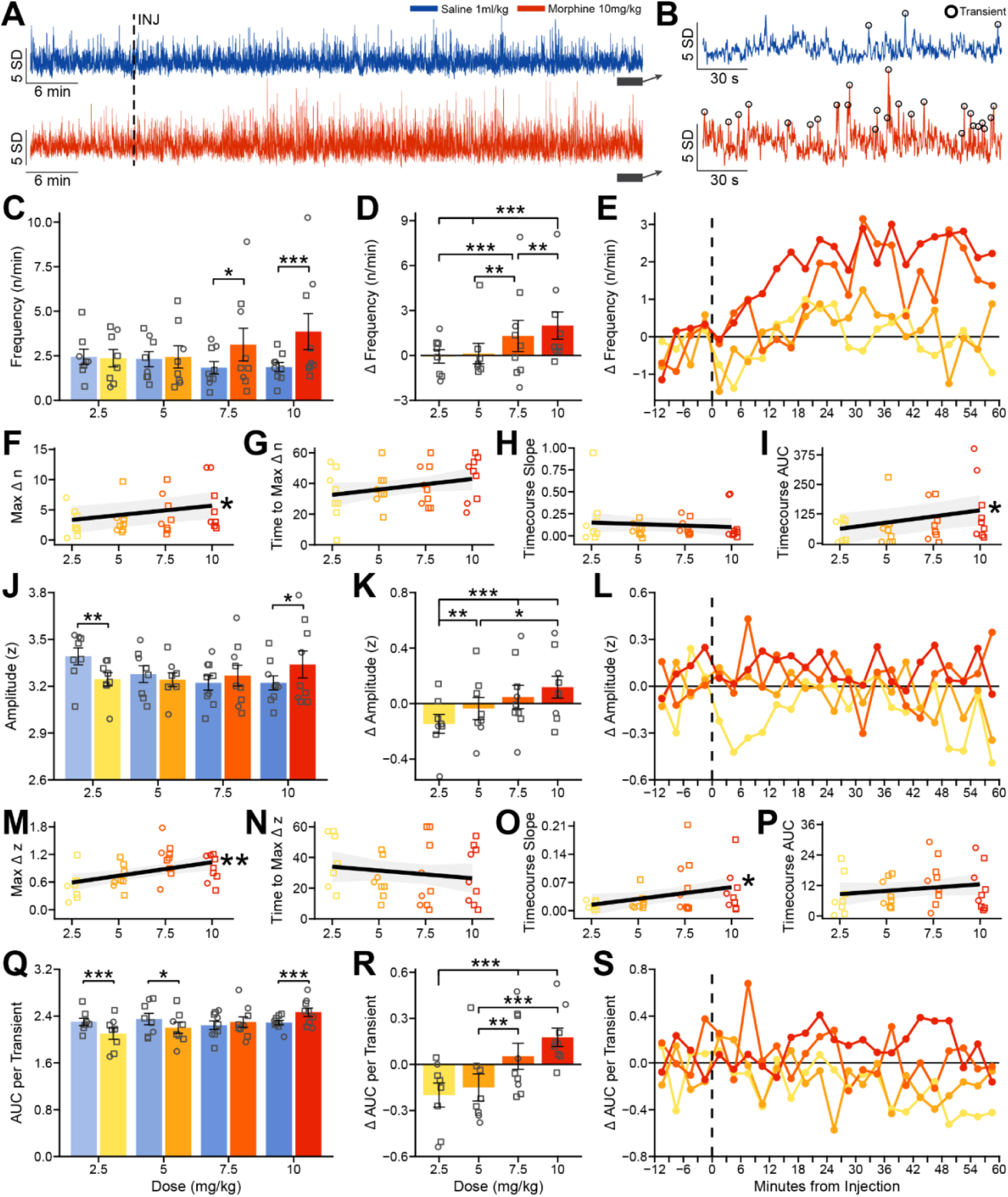
Morphine treatment increases VTA DA GCaMP6f transient frequency, amplitude, and area under the curve in a dose-dependent manner. **A)** Representative whole session traces of the 10.0 mg/kg morphine dose condition for saline control (top) and morphine treatment (bottom) sessions. Whole session streams were normalized (Z Score) to the mean and standard deviation of the within session pre-injection baseline. **B)** Example of spontaneous transient detection (30s) from the traces A. Circles mark identified transients. **C)** Mean transient frequency (events per minute) by injection type (saline or morphine) and dose (2.5, 5.0, 7.5, and 10.0 mg/kg). Individual points represent each subject (male = square, female = circle). Cool colors (blues) indicate saline control sessions and warm colors (oranges) indicate morphine sessions. **D)** Change in transient frequency with morphine treatment relative to the matched saline control session by dose. **E)** Change in transient frequency with morphine treatment by dose relative to the matched saline control session across the session (3-minute time bins). Colors correspond to doses in C, D. Time course analysis on change in transient frequency relative to matched saline control sessions across increasing doses of morphine: **F)** Maximum change in transient frequency, **G)** Time to the maximum change in frequency, **H)** Slope of the timecourse from injection to the maximum change in frequency, and **I)** Cumulative area under the curve of the timecourse. **J)** Mean transient amplitude (Z score) by injection type and dose. **K)** Morphine treatment change in transient amplitude relative to the matched saline control session by dose. **L)** Change in transient amplitude with morphine treatment by dose relative to the matched saline control session across 3-minute time bins. Colors correspond to doses in J, K. Time course analysis on change in transient amplitude relative to matched saline control sessions across increasing doses of morphine: **M)** Maximum change in amplitude, **N)** Time to the maximum change in amplitude, **O)** Slope of the timecourse from injection to the maximum change in amplitude, and **P)** Cumulative area under the curve of the transient amplitude timecourse. **Q)** Mean transient area under the curve (AUC) by injection type and dose. **R)** Change in transient AUC relative to matched saline control sessions across morphine doses. **S)** Change in transient AUC with morphine treatment by dose across 3-minute time bins. Colors correspond to doses in Q, R. * <0.05; **<0.01; ***<0.001

##### Morphine session relative to saline control

To characterize morphine-evoked changes for each treatment session, we computed the difference (Δ) between each morphine session and the prior day’s saline session. For each outcome (frequency, amplitude, AUC), the change value (M-S) was calculated for each time bin. For amplitude and AUC, difference was computed from average values in each time bin between sessions. Separate mixed effects models were fit to the change values with fixed effects for dose and time bin and the interaction between them. Results averaging across the whole session are shown in Figures 2, 3, and 4, panels D, K, and R. The bin-by-bin timecourses of the difference between morphine and saline sessions are shown in Figures 2, 3, and 4, panels E, L, and S.

##### Timecourse of transient events

To further assess temporal dynamics of morphine treatment effects, we estimated the rate of change of transient characteristics following injection. For both transient frequency and amplitude at each morphine dose, we calculated the pre-injection Baseline mean (mean of all time bins before the injection), maximum change (Max Δ; maximum post-injection time bin change value) and time to maximum change (Time to Max Δ; minutes post-injection of the maximum post-injection time bin change value). We then fit the Timecourse Slope with linear regression from Baseline at time 0 (injection) through Max Δ at Time to Max Δ. Results from these analyses are shown in Figures 2, 3, and 4, panels F-H and M-O. To summarize the cumulative change in transient frequency and amplitude across the session, we calculated Timecourse AUC using trapezoidal numerical integration after subtracting the baseline and rectifying to retain only positive excursions above baseline. Results from these analyses are shown in Figures 2, 3, and 4, panels I and P.

Dose-dependent effects of the magnitude of morphine-evoked changes in transient parameters were modeled using linear mixed effects models separately for Max Δ, Time to Max Δ, Timecourse Slope, and Timecourse AUC. Models included a fixed effect of dose (continuous) and a random intercept by subject. Model fit was characterized with conditional R^2^ values (Nakagawa & Schielzeth, 2013). Predicted values across the observed dose range were generated with 95% confidence intervals.

##### Sensor transient comparison

To compare the magnitude of changes in spontaneous events across morphine doses between sensors, we computed change scores for each subject, dose, and time bin. We first paired each morphine treatment session with the corresponding saline control session by subject and time bin. For every transient event variable (frequency, amplitude, AUC), the difference was calculated for each time bin as:

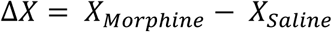

Change values were standardized using the pooled baseline variability within each sensor type to allow for comparison across sensors and outcome scales. Specifically, for each sensor we computed the standard deviation of saline-only sessions and used these values to scale the morphine-saline difference scores as:

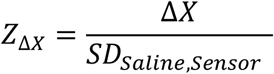

This normalization procedure yields a signal-to-noise-like standardized change metric that reflects the magnitude of morphine-induced changes in spontaneous transients relative to baseline variability.

To assess morphine-induced changes across sensors, we implemented linear mixed-effects models in glmmTMB (Brooks et al., 2017) for each outcome variable. Analyses were restricted to post-injection time bins of the morphine sessions. First, a base model was fit with fixed effects and interactions of dose (centered; continuous), sensor, and time bin. A random intercept by subject was included to account for repeated measurements within subjects. A second model was fit with the AR(1) structure and used to evaluate fixed-effect terms when it provided a lower AIC than the base model. Type II Wald Χ^2^ tests (Fox & Weisberg, 2019) were used to evaluate the significance of fixed effects. Estimate marginal means, joint tests, pairwise comparisons, and dose-response slopes were computed to characterize sensor differences at each dose level.

##### Whole session sensor comparison

To compare changes in whole stream mean and MAD between sensors, we computed change scores for each subject, dose, and time bin relative to the corresponding saline control session. Change values were assessed between sensors via linear mixed-effects models in glmmTMB as described for between sensor transient change models. Estimate marginal means, joint tests, pairwise comparisons, and dose-response slopes were computed to characterize sensor differences in whole session mean and MAD at each dose level.

## 3. RESULTS

### 3.1. Baseline Measurement of Dopamine Transients at Each Sensor

For each sensor, we first characterized spontaneous transients during the first saline control session (Fig. 1F). Transient frequency and amplitude was highest with the dLight1.3b sensor compared with GCaMP6f and GRABDA2h (Frequency: GCaMP6f vs dLight1.3b *t*(572) = -1.68, *p =* 0.001, *d* = -1.41, dLight1.3b vs GRABDA2h *t*(572) = 2.91, *p* < 0.0001, *d* = 2.45, Fig. 1G; Amplitude: GCaMP6f vs dLight1.3b *t*(528) = -0.42, *p* < 0.0001, *d* = -1.06, dLight1.3b vs GRABDA2h *t*(528) = 0.62, *p* < 0.0001, *d* = 1.58, Fig. 1H). Transient frequency and amplitude with GCaMP6f was higher than with GRABDA2h (Frequency: *t*(572) = 1.23, *p =* 0.022, *d* = 1.04, Fig. 1G; Amplitude: *t*(528) = 0.20, *p =* 0.035, *d* = 0.52, Fig. 1H). Likewise, average AUC was largest for dLight1.3b transients compared with GCaMP6f and GRABDA2h (GCaMP6f vs dLight1.3b: *t*(528) = -0.48, *p =* 0.014, d = -0.65; dLight1.3b vs GRABDA2h: *t*(528) = 1.37, *p* < 0.0001, d = 1.83; Fig. 1I), with GRABDA2h also having less AUC per transient than GCaMP6f (*t*(528) = 0.88, *p* < 0.0001, d = 1.18). Differences in AUC can be explained by differences in amplitude but also in kinetics. Transients from dLight1.3b had the shortest rise duration, corresponding to a steeper slope (Rise Duration: GCaMP6f vs dLight1.3b *t*(526) = 0.57, *p* < 0.0001, d = 0.64, dLight1.3b vs GRABDA2h *t*(526) = -0.48, *p =* 0.0005, d = -0.54, Fig. 1J; Rise Slope: GCaMP6f vs dLight1.3b *t*(528) = -0.005, *p* < 0.0001, d = -2.04, dLight1.3b vs GRABDA2h *t*(528) = 0.004, *p* < 0.0001, d = 1.50, GCaMP6f vs GRABDA2h *t*(528) = -0.001, *p =* 0.025, d = -0.54, Fig. 1K). In addition, the falling phase of transients depended on sensor, with dLight1.3b showing the shortest duration of the falling phase (Fig. 1L) and steepest negative slope (Fig. 1M). GRABDA2h had the longest duration and most shallow slope, with GCaMP6f falling intermediate between the two (Fall Duration: GCaMP6f vs dLight1.3b *t*(528) = 0.50, *p =* 0.055, d = 0.86, dLight1.3b vs GRABDA2h *t*(528) = -1,72, *p* < 0.0001, d = -2.98, GCaMP6f vs GRABDA2h *t*(528) = -1.22, *p* < 0.0001, d = -2.12; Fall Slope: GCaMP6f vs dLight1.3b *t*(527) = 1.33, *p* < .0001, d = 2.50, dLight1.3b vs GRABDA2h *t*(527) = -1.74, *p* < 0.0001, d = -3.26, GCaMP6f vs GRABDA2h *t*(527) = - 0.41, *p =* 0.053, d = -0.76).

### 3.2. VTA_DA_ GCaMP6f Transients

To assess the effects of acute morphine administration on GCaMP6f signaling in VTA_DA_ cell bodies, we tested four morphine doses in ascending order (2.5, 5, 7.5, and 10 mg/kg, i.p.), each preceded by a volume-matched saline control session on the prior day (Fig. 1D). Each session consisted of a 12-minute baseline recording period prior to injection followed by 60-minutes of recording post-injection (Fig. 1E). Representative traces of saline control and morphine treatment sessions (10 mg/kg) and transient detection are shown in Fig. 2A and Fig. 2B respectively.

#### 3.2.1. Transient Frequency

##### Overall Effects of Morphine

Morphine administration exerted a significant dose-dependent effect on spontaneous transient frequency. There was a main effect of dose (Χ^2^ = 11.35, *p =* 0.010) as well as a significant interaction between treatment (saline versus morphine) and dose (Χ^2^ = 64.80, *p* < 0.0001). There was no main effect of time bin (Χ^2^ = 20.61, *p =* 0.359) or interactions with other factors (all *p* > 0.20).

##### Morphine Effects on Transient Frequency

Morphine administration increased overall session transient frequency (n/min) at VTA_DA_ GCaMP6f sensors in a dose-dependent manner (Fig. 2C). At 2.5 and 5 mg/kg, morphine did not significantly alter frequency compared to matched saline controls (2.5 mg/kg: M_S_ = 2.44, M_M_ = 2.37; *t*(1198) = -0.24, *p =* 0.672, *d* = -0.15; 5 mg/kg: M_S_ = 2.31, M_M_ = 2.44; *t*(1198) = -0.80, *p =* 0.424, *d* = -0.29). At the 7.5 mg/kg dose, transient frequency significantly increased from 1.83 transients/min in the saline condition to 3.13 with morphine (*t*(1198) = -2.26, *p =* 0.024, *d* = -0.82). At 10 mg/kg, transient frequency increased from 1.87 to 3.86 (*t*(1198) = -3.48, *p =* 0.0005, d = -1.26).

Because there was a decrease in transient frequency in saline control sessions over the course of the experiment (*F*(3,1198) = 8.42, *p* < 0.0001; Fig. 2C), we calculated the overall average change in transient frequency for each morphine session relative to its paired saline session. The change (Δ) in frequency scaled with morphine dose (Χ^2^ = 61.31, *p* < 0.0001; Fig. 2D). The highest dose of 10 mg/kg drove the greatest increase in transient frequency relative to all lower doses: 2.5 mg/kg (*t*(598) = -6.92, *p* < 0.001, d = - 0.76), 5 mg/kg (*t*(598) = -6.05, *p* < 0.001, d = -0.66), and 7.5 mg/kg (*t*(598) = -2.86, *p =* 0.004, d = -0.30). Administration of 7.5 mg/kg morphine drove greater increases relative to 2.5 mg/kg (*t*(598) = -4.17, *p* < 0.001, d = -0.46) and 5 mg/kg (*t*(598) = -3.30, *p =* 0.001, d = -0.36). No difference was observed between 2.5 and 5 mg/kg (*t*(598) = -0.83, *p =* 0.405, d = -0.10).

##### Timecourse of Transient Frequency

To characterize the pattern of morphine effects on transient frequency in VTA_DA_ cell bodies over the course of the session, we analyzed bin-by-bin change in transient frequency of each morphine session from its matched saline control. During the 60-minute treatment period, the average elevation of transient frequency in morphine sessions increased across time bins (Fig. 2E). From this timecourse, we estimated the maximum change in frequency (Max Δn), Time to Max Δn, the slope of the timecourse from injection to Max Δn, and cumulative AUC of the timecourse. The maximum change in frequency depended on dose (β = 0.31, SE = 0.13, *p =* 0.017; Fig. 2F), with increases in transient frequency during morphine sessions ranging from 2.79 transients/min at the lowest dose to 5.41 at the highest. Time to reach Max Δn, though shorter for 2.5 mg/kg (32.63 min) than 10 mg/kg (43.67 min), did not significantly vary by dose, and was on average 37.76 minutes (β = 1.38, SE = 0.81, *p =* 0.092; Fig. 2G). The slope of the timecourse change in frequency did not depend on dose (β = -0.006, SE = 0.01, *p =* 0.529; Fig. 2H), but cumulative AUC of the timecourse did (β = 10.43, SE = 4.18, *p =* 0.012; Fig. 2I). The dose-dependent increase in timecourse AUC reflects that transient frequency showed overall greater increases with higher doses of morphine.

#### 3.2.2. Transient Amplitude

##### Overall Effects of Morphine

In addition to increasing the frequency of spontaneous transients in VTA_DA_ cell bodies, morphine administration modulated the amplitude of spontaneous events. While there was no significant main effect of treatment (Χ^2^ = 0.03, *p =* 0.854), we observed a significant main effect of dose (Χ^2^ = 7.83, *p =* 0.0497) and a significant interaction between treatment and dose (Χ^2^ = 52.38, *p* < 0.0001). No additional interactions were significant (all *p* > 0.05).

##### Morphine Effects on Transient Amplitude

Morphine administration exerted a bidirectional dose-dependent effect on overall session transient amplitude (Fig. 2J). At the lowest dose of 2.5 mg/kg, morphine administration significantly reduced transient amplitude relative to the matched saline control session (M_S_ = 3.39, M_M_ = 3.24; *t*(10195) = -2.79, *p =* 0.005, *d* = 0.23). Intermediate doses (5 and 7.5 mg/kg) produced no significant deviation from the control session (5 mg/kg: M_S_ = 3.28, M_M_ = 3.24; *t*(10195) = 0.90, *p =* 0.367, *d* = 0.07; 7.5 mg/kg: M_S_ = 3.22, M_M_ = 3.27; *t*(10195) = -0.98, *p =* 0.326, *d* = 0.08). The highest dose of 10 mg/kg produced a significant increase in transient amplitude compared with the matched control session (M_S_ = 3.22, M_M_ = 3.34; *t*(10195) = -2.41, *p =* 0.016, *d* = -0.19).

Because transient amplitude decreased across saline control sessions (*F*(3,10195) = 12.04 *p* < 0.0001; Fig. 2J), we calculated the change in amplitude for each morphine dose from its paired saline session. The change in amplitude depended on morphine dose (Χ^2^ = 34.84, *p* < 0.0001; Fig. 2K). Relative to 2.5 mg/kg, the 7.5 and 10 mg/kg doses showed increases in transient amplitude (10 mg/kg: (*t*(539) = -5.64, *p* < 0.0001, d = -0.645; 7.5 mg/kg (*t*(539) = -4.66, *p* < 0.0001, d = -0.54; 5 mg/kg: (*t*(539) = - 2.92, *p =* 0.004, d = -0.35). Change in transient amplitude was also greater with 10 mg/kg relative to 5 mg/kg (*t*(539) = -2.54, *p =* 0.012, d = -0.29). No differences were observed between 5 and 7.5 mg/kg (*t*(539) = -1.63, *p =* 0.105, d = -0.19) or 7.5 and 10 mg/kg (*t*(539) = -0.90, *p =* 0.368, d = -0.10).

##### Timecourse of Transient Amplitude

As with frequency, we analyzed the change in transient amplitude between each morphine session and its matched saline control session across time bins (Fig. 2L). The maximum change in amplitude (Max Δz) across time bins ranged from 0.53 to 0.90, significantly depending on dose (β = 0.06, SE = 0.02, *p =* 0.003; Fig. 2M). The time to Max Δz did not depend on dose, with maximum change in amplitude reached at 30.19 minutes on average (β = -1.01, SE = 1.06, *p =* 0.342; Fig. 2N). The slope of the timecourse increased significantly across doses, reflecting the increase in amplitude in the same timeframe (β = 0.01, SE = 0.003, *p =* 0.040; Fig. 2O), but the cumulative AUC of the timecourse of transient amplitude, while positive, did not depend on dose (β = 0.51, SE = 0.47, *p =* 0.272; Fig. 2P). The increase in transient amplitude reached higher levels with higher doses of morphine, but, like frequency, did not take longer to reach the maximum level.

#### 3.2.3. Transient Area Under the Curve

##### Overall Effects of Morphine

To identify changes in the shape of transient events in VTA_DA_ cell bodies, AUC was calculated for each transient event as total area under the curve relative to pre-event peak baseline. Transient AUC depended on dose (Χ^2^ = 55.53, *p* < 0.0001) and time bin (Χ^2^ = 35.83, *p =* 0.011), as well as a significant interaction between treatment and dose (Χ^2^ = 85.95, *p* < 0.0001). No significant main effect of treatment (Χ^2^ = 0.06, *p =* 0.802) or interactions between other fixed effects were significant (all *p* > 0.07).

##### Morphine Effects on Transient AUC

The effects of morphine on transient AUC were similar to those on amplitude (Fig. 2Q). At lower doses, AUC was lower for morphine sessions compared to matched saline control, with significant decreases observed at 2.5 mg/kg (M_S_ 2.30, M_M_ = 2.10; *t*(10194) = 3.63, *p =* 0.0003, *d* = 0.24) and 5 mg/kg (M_S_ = 2.35, M_M_ = 2.20; *t*(10194) = 2.57, *p =* 0.010, *d* = 0.17). No effect was observed at the 7.5 mg/kg dose (M_S_ = 2.24, M_M_ = 2.30; *t*(10194) = -1.54, *p =* 0.124, *d* = -0.10). At 10 mg/kg, AUC was significantly increased by morphine administration (M_S_ = 2.29, M_M_ = 2.46; *t*(10194) = -3.72, *p =* 0.0002, *d* = -0.24).

Although there was no consistent trend across saline control sessions, we calculated change in AUC of morphine sessions compared with their paired saline sessions to match analyses of frequency and amplitude. We observed a significant effect of dose on the change in AUC (Χ^2^ = 46.07, *p* < 0.0001; Fig. 2R). Transient AUC was reduced with lower doses of morphine relative to saline control, with no difference in the change in transient AUC between 2.5 and 5 mg/kg (*t*(539) = -1.32, *p =* 0.186, d = -0.16). Both higher morphine doses increased AUC relative to saline control, with no difference between 7.5 and 10 mg/kg (*t*(539) = -1.60, *p =* 0.110, d = -0.18). The change in AUC was higher for 10 mg/kg compared to 2.5 mg (*t*(539) = -6.06, *p* < 0.0001, d = -0.69) and 5 mg/kg (*t*(539) = -4.63, *p* < 0.0001, d = -0.53). The 7.5 mg/kg dose also significantly increased the change in AUC relative to the 2.5 mg/kg (*t*(539) = -4.41, *p* < 0.0001, d = - 0.51) and 5 mg/kg (*t*(539) = -3.01, *p =* 0.003, d = -0.35) doses. No main effect of time bin (Χ^2^ = 24.50, *p =* 0.178) or interaction between time bin and dose (Χ^2^ = 66.84, *p =* 0.175) were observed on the change in AUC (Fig. 2S).

Overall, we observed dose-related increases in transient event frequency, amplitude, and AUC following morphine treatment via the GCaMP6f sensor, indicating increased VTA_DA_ cell body activity.

### 3.3. NAcLS dLight1.3b Transients

To compare downstream DA release with VTA_DA_ activity, we assessed the effects of ascending morphine doses on dLight1.3b activity in NAcLS. Overall, morphine administration significantly modulated spontaneous DA transients. Representative traces of saline control and morphine treatment sessions (10 mg/kg) and transient detection are shown in Fig. 3A and Fig. 3B. All analyses follow the same conventions as Figure 2.

**Figure 3.**
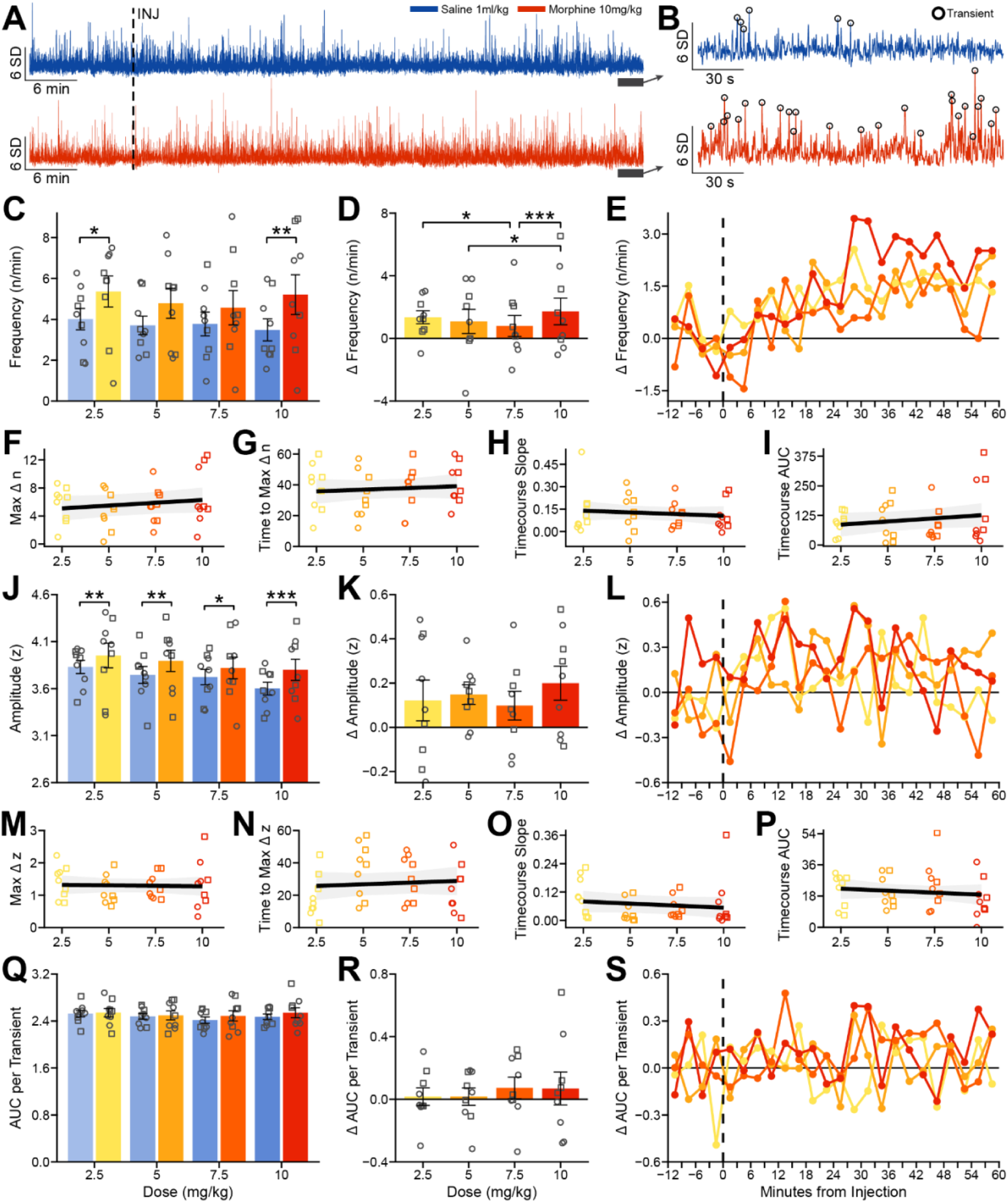
Morphine treatment increases NAcLS dLight1.3b transient frequency and amplitude. Same conventions as Figure 2.

#### 3.3.1. Transient Frequency

##### Overall Effects of Morphine

Morphine administration had a significant dose-dependent effect on spontaneous transient event frequency. We observed significant main effects of treatment (Χ^2^ = 4.03, *p =* 0.045) and dose (Χ^2^ = 17.48, *p* < 0.0001), and a significant interaction between treatment and dose (Χ^2^ = 13.26, *p =* 0.004). No main effect of time bin was observed (Χ^2^ = 27.72, *p =* 0.089) and time bin did not interact with other factors (all *p* > 0.10).

##### Morphine Effects on Transient Frequency

Morphine administration increased the frequency of dLight1.3b transients in NAcLS (Fig. 3C). Increases in transient frequency for morphine sessions relative to matched saline controls were significant at 2.5 mg/kg (M_S_ = 4.01, M_M_ = 5.36; *t*(1276) = -2.40, *p =* 0.017, *d* = -0.75) and 10 mg/kg (M_S_ = 3.48, M_M_ = 5.21; *t*(1276) = -3.07, *p =* 0.002, *d* = -0.96). Intermediate doses did not produce a significant increase in transient frequency (5 mg/kg: M_S_ = 3.70, M_M_ = 4.57; *t*(1276) = - 1.92, *p =* 0.0546, *d* = -0.61; 7.5 mg/kg: M_S_ = 3.77, M_M_ = 4.57; *t*(1276) = -1.41, *p =* 0.159, *d* = -0.44).

As transient frequency across saline control sessions decreased over time (*F*(3, 1276) = 2.68, *p =* 0.046; Fig. 3C), we compared each morphine session to its matched saline control session. The change in transient frequency from matched saline control sessions was significantly affected by morphine dose (Χ^2^ = 13.25, *p =* 0.004; Fig. 3D). The 10 mg/kg dose resulted in greater increases in the change in transient frequency relative to the 5 mg/kg (*t*(636) = -2.41, *p =* 0.016, d = -0.25) and 7.5 mg/kg (*t*(636) = -3.49, *p* < 0.001, d = -0.37) doses. The change in transient frequency of the 2.5 mg/kg dose was also higher than the 7.5 mg/kg dose (*t*(636) = 2.08, *p =* 0.038, d = 0.22). No differences were observed between 2.5 and 5 mg/kg (*t*(636) = 1.00, *p =* 0.318, d = 0.11), 2.5 and 10 mg/kg (*t*(636) = -1.41, *p =* 0.159, d = -0.15), and 5 and 7.5 mg/kg (*t*(636) = 1.08, *p =* 0.279, d = 0.11).

##### Timecourse of Transient Frequency

Across the session, the average increase in transient frequency following morphine treatment, relative to saline control session significantly depended on time (Χ^2^ = 30.09, *p =* 0.050; Fig. 3E) but there were no interactions between dose and time bin (Χ^2^ = 33.26, *p =* 0.995). Across time bins, the maximum increase in transient frequency by morphine did not depend on dose (β = 0.16, SE = 0.13, *p =* 0.200; Fig. 3F), with a mean maximum increase from saline of 5.69 transients/min. The mean time to maximum increase was 37.32 minutes, which was not dose-dependent (β = 0.44, SE = 0.75, *p =* 0.556; Fig. 3G). The slope of the timecourse to maximum change in frequency did not depend on morphine dose (β = -0.005, SE = 0.01, *p =* 0.490; Fig. 3H), nor did cumulative AUC of the timecourse (β = 5.31, SE = 4.18, *p =* 0.203; Fig. 3I). Thus, while acute morphine treatment increased the frequency of dLights1.3b transients in NAcLS, no differences in the timecourse of morphine effects were observed across doses.

#### 3.3.2. Transient Amplitude

##### Overall Effects of Morphine

Morphine treatment modulated the amplitude of dLight1.3b transients in NAcLS. Transient amplitude was affected by treatment (Χ^2^ = 17.69, *p* < 0.0001), dose (Χ^2^ = 43.60, *p* < 0.0001), and time bin (Χ^2^ = 42.76, *p =* 0.001), as well as significant interactions between treatment and time bin (Χ^2^ = 39.33, *p =* 0.004) and dose and time bin (Χ^2^ = 86.46, *p =* 0.007).

##### Morphine Effects on Transient Amplitude

Morphine administration increased transient amplitude relative to matched saline control sessions (Fig. 3J). The observed increase in amplitude was significant at 2.5 mg/kg (M_S_ = 3.83, M_M_ = 3.95; t(18679) = -3.26, *p =* 0.001, d = -0.12), 5 mg/kg (M_S_ = 3.75, M_M_ = 3.89; t(18679) = -3.19, *p =* 0.001, d = -0.12), 7.5 mg/kg (M_S_ = 3.72, M_M_ = 3.82; t(18679) = -1.96, *p =* 0.0502, d = -0.08), and 10 mg/kg (M_S_ = 3.60, M_M_ = 3.80; t(18679) = -3.93, *p =* 0.0001, d = -0.15). Across saline control sessions, transient amplitude decreased across experimental sessions (F(3,18679) = 7.55, *p* < 0.0001). While all morphine doses produced an overall increase in transient amplitude relative to matched saline control sessions, no significant differences between morphine doses were observed (Χ^2^ = 4.16, *p =* 0.244; Fig. 3K).

##### Timecourse of Transient Amplitude

The change in dLight1.3b transient amplitude significantly depended on time (Χ^2^ = 33.73, *p =* 0.020; Fig. 3L) but there was no interaction between dose and time bin (Χ^2^ = 65.80, *p =* 0.198). Over the timecourse, the maximum change in amplitude was a consistent increase of 1.30 z-score, with no significant differences between doses in the maximum increase (β = -0.006, SE = 0.03, *p =* 0.836; Fig. 3M). Time to maximum increase in amplitude also was consistent across doses with an average of 27.25 minutes (β = 0.41, SE = 0.79, *p =* 0.599; Fig. 3N). The slope of the timecourse (β = -0.003, SE = 0.004, *p =* 0.330; Fig. 3O) and cumulative AUC of the timecourse of transient amplitude (β = -0.46, SE = 0.64, *p =* 0.468; Fig. 3P) also were not different between doses. Note that although there were not dose-dependent differences in the increase in amplitude detected at dLight1.3b sensors, the time to reach the maximum change largely aligned with those observed with GCaMP6f.

#### 3.3.3. Transient Area Under the Curve

##### Overall Effects of Morphine

Transient shape was quantified as AUC for each event. The average AUC of dLight1.3b transients depended on dose (Χ^2^ = 12.75, *p =* 0.005) and time bin (Χ^2^ = 56.45, *p* < 0.0001) as well as interactions between dose and time bin (Χ^2^ = 124.87, *p* < 0.0001) and treatment, dose, and time bin (Χ^2^ = 93.07, *p =* 0.002). From the interactions, post-hoc tests did not reveal any consistent pattern of effects. No significant main effect of treatment (Χ^2^ = 1.35, *p =* 0.246) or interactions between other fixed effects were significant (all *p* > 0.06).

##### Morphine Effects on Transient AUC

Although there was an overall significant effect of dose, there were no significant differences between the individual saline and morphine sessions (all *p* > 0.05; Fig. 3Q). There were no changes across saline control sessions (F(3,18669) = 2.40, *p =* 0.066). Similarly, the change in AUC in morphine sessions compared with their paired saline sessions did not vary with dose (Χ^2^ = 3.78, *p =* 0.286; Fig. 3R). There was no main effect of time bin (Χ^2^ = 22.92, *p =* 0.241) nor interaction between time bin and dose (Χ^2^ = 69.02, *p =* 0.132) on the change in AUC by morphine as well (Fig. 3S).

Overall, we observed increases in the frequency and amplitude of transient events from the dLight1.3b sensor, indicating increased release of DA in NAcLS following morphine treatment. While increases in transient frequency and amplitude were consistently observed, differentiation between doses was not as clear relative to those observed with VTA_DA_ GCaMP6f.

### 3.4. NAcLS GRABDA2h Transients

GRABDA2h is another dopamine sensor but is modified from the D2 dopamine receptor. We measured the effects of ascending doses of morphine on the activity of GRABDA2h. Representative traces of saline control and morphine treatment sessions (10 mg/kg) and transient detection are shown in Figs. 4A and B. Overall, effects on NAcLS GRADBA2h spontaneous transients were inconsistent with observations with VTA_DA_ GCaMP6f and NAcLS dLight1.3b. Analyses that follow use the same conventions as Figures 2 and 3.

**Figure 4.**
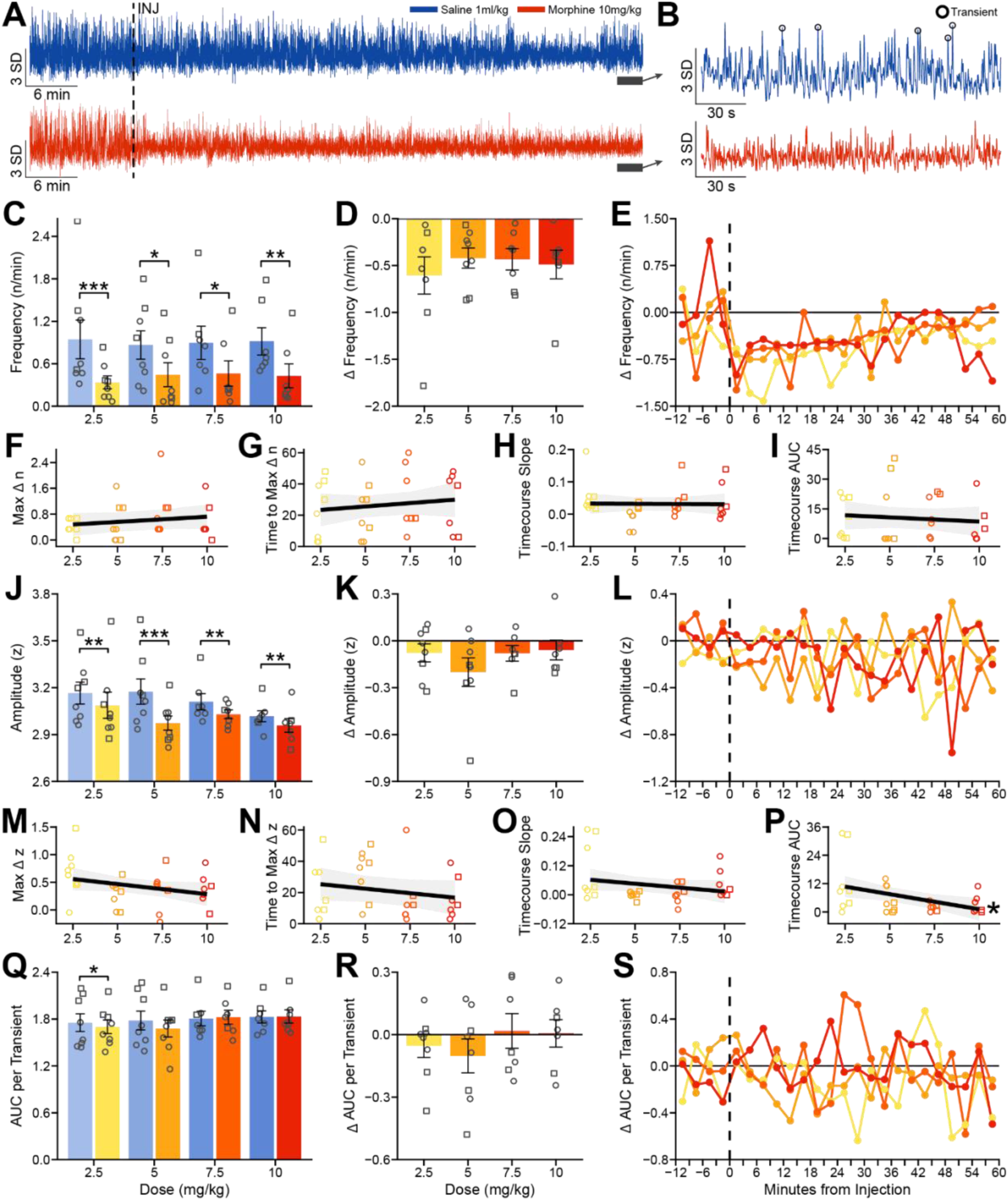
Morphine treatment decreases NAcLS GRABDA2h transient frequency and amplitude in a dose-independent manner. Same conventions as Figure 2.

#### 3.4.1. Transient Frequency

##### Overall Effects of Morphine Administration

Morphine administration significantly affected spontaneous transient events. Transient frequency from GRABDA2h sensors depended on treatment (Χ^2^ = 10.58, *p =* 0.001) and time (Χ^2^ = 90.96, *p* < 0.0001). No significant main effect of dose (Χ^2^ = 0.16, *p =* 0.983) or interactions between other variables were observed (all *p* > 0.10).

##### Morphine Effects on Transient Frequency

Unlike GCaMP6f and dLight1.3b, the frequency of GRABDA2h transients following morphine administration decreased relative to matched saline control sessions across all doses (Fig. 4C). Decreases in transient frequency relative to saline control were significant with 2.5 mg/kg (M_S_ = 0.94, M_M_ = 0.34; t(1036) = 3.51, *p* < 0.001, d = 1.14), 5 mg/kg (M_S_ = 0.86, M_M_ = 0.44; t(1036) = 2.43, *p =* 0.015, d = 0.79), 7.5 mg/kg (M_S_ = 0.90, M_M_ = 0.46; t(1036) = 2.46 *p =* 0.014, d = 0.81), and 10 mg/kg doses (M_S_ = 0.92, M_M_ = 0.43; t(1036) = 2.77, *p =* 0.006, d = 0.91).

Saline control sessions were stable across doses, with no significant differences observed (F(3,1036) = 0.70, *p =* 0.552). Across morphine doses, the changes in transient frequency relative to matched saline control did not differ from each other (Χ^2^ = 6.84, *p =* 0.077; Fig. 4D).

##### Timecourse of Transient Frequency

For each dose of morphine relative to its control saline session, there were no shifts in the timecourse of change in responses across doses (Fig. 4E). No significant main effect of time bin (Χ^2^ = 26.98, *p =* 0.105) or interaction between dose and time bin (Χ^2^ = 66.96, *p =* 0.172) were observed. The mean maximum change in transient frequency was 0.61/min, with no dependence on dose (β = 0.032, SE = 0.03, *p =* 0.288; Fig. 4F). Time to reach the maximum increase was on average 26.71 minutes, and also did not depend on dose (β = 0.87, SE = 1.17, *p =* 0.455; Fig. 4G).The slope of the timecourse did not vary with dose (β = -0.0003, SE = 0.003, *p =* 0.925; Fig. 4H), nor did cumulative AUC of the timecourse (β = -0.43, SE = 0.73, *p =* 0.562; Fig. 4I).

#### 3.4.2. Transient Amplitude

##### Overall Effects of Morphine

Morphine treatment also modulated the amplitude of GRABDA2h transients in NAcLS. Treatment (Χ^2^ = 39.66, *p* < 0.0001), dose (Χ^2^ = 19.98, *p =* 0.0002), and time bin (Χ^2^ = 61.80, *p* < 0.0001) had significant main effects on amplitude. No significant interactions were observed.

##### Morphine Effects on Transient Amplitude

Morphine administration decreased transient amplitude relative to the matched saline control sessions (Fig. 4J). The observed decrease in amplitude from saline control was significant with all morphine doses (2.5 mg/kg: M_S_ = 3.17, M_M_ = 3.09; t(2216) = 2.81, *p =* 0.005, d = 0.29; 5 mg/kg: M_S_ = 3.17, M_M_ = 2.97; t(2216) = 3.79, *p =* 0.0002, d = 0.35; 7.5 mg/kg: M_S_ = 3.11, M_M_ = 3.03; t(2216) = 2.96, *p =* 0.003, d = 0.30; 10 mg/kg: M_S_ = 3.02, M_M_ = 2.96; t(2216) = 2.82, *p =* 0.005, d = 0.29).

Across saline control sessions, there was a significant decrease in transient amplitude (F(3,2216) = 3.40, *p =* 0.017). When comparing transient amplitude from each morphine session to its matched saline sessions, there was no effect of dose on the reduction in transient amplitude across morphine doses (Χ^2^ = 2.65, *p =* 0.449; Fig. 4K). **Timecourse of Transient Amplitude.** The change in GRABDA2h transient amplitude in each morphine sessions relative to its saline control did not depend on time (Χ^2^ = 13.75, *p =* 0.798) or an interaction between dose and time bin (Χ^2^ = 56.65, *p =* 0.488; Fig. 4L). Over the course of the sessions, the maximum change in amplitude did not differ (average = 0.42 z-score) and was not affected by dose (β = -0.04, SE = 0.02, *p =* 0.104; Fig. 4M). Time to maximum change in amplitude also did not depend on dose (β = -1.15, SE = 0.96, *p =* 0.231; Fig. 4N) and was 21.28 minutes on average. The timecourse slope also did not vary by dose (β = -0.006, SE = 0.005, *p =* 0.199; Fig. 4O). The only significant relationship was a negative effect of dose on the cumulative AUC of the timecourse of transient amplitude relative to saline control sessions across time (β = -1.29, SE = 0.50, *p =* 0.010; Fig. 4P).

#### 3.4.3. Transient Area Under the Curve

##### Overall Effects of Morphine

AUC for individual transient events depended on dose (Χ^2^ = 11.73, *p =* 0.008). No other significant main effects or interactions were observed (all *p* > 0.06).

##### Morphine Effects on Transient AUC

At the 2.5 mg/kg dose, there was a decrease in average AUC of individual transients following morphine administration relative to the matched saline control (t(2215) = 2.07, *p =* 0.039, d = 0.21; Fig. 4Q). No significant differences in transient AUC relative to saline control were observed with 5, 7.5, or 10 mg/kg doses (all *p* > 0.15). Transient AUC was consistent across saline control sessions (F(3, 2215) = 0.89, *p =* 0.446). Across morphine treatment doses, no effect of dose on the change in AUC relative to matched saline session was observed (Χ^2^ = 3.73, *p =* 0.292; Fig. 4R). The difference in AUC in morphine sessions relative to matched saline sessions also did not significantly vary as a function of time bin (Χ^2^ = 19.26, *p =* 0.440) or interaction between time bin and dose (Χ^2^ = 52.75, *p =* 0.635; Fig. 4S).

Overall, we observed patterns of decreases in transient events from the GRABDA2h sensor. On its face, such findings would indicate reduced DA signaling in NAcLS, in contrast to observations from signals from dLight1.3b sensors in the same region following morphine treatment. Further, these findings contrast with observed activity of VTA_DA_ cells, as assayed by GCaMP6f signal.

### 3.5. Between Sensor Transient Comparison

Qualitatively, the patterns of responses to morphine differed across GCaMP6f, dLight1.3b, and GRABDA2h sensors. To make quantitative comparisons of the effects of morphine on DA activity across sensors, we normalized each metric (frequency, amplitude, and transient AUC) for each sensor to allow for comparison between sensors despite absolute differences in transient frequency and kinetics. Figure 5A-C show overlaid individual transient events from one representative subject for the effects of the 10 mg/kg dose on signaling at the VTA_DA_ GCaMP6f (Fig. 5A), NAcLS dLight1.3b (Fig. 5B), and NAcLS GRABDA2h (Fig. 5C) sensors. In these panels, color indicates the density of overlap of individual traces. Note that the scale of the colormap indicating number of traces varies across sensors, reflecting differences in overall frequency of transients, consistent with Figure 1F-G.

**Figure 5.**
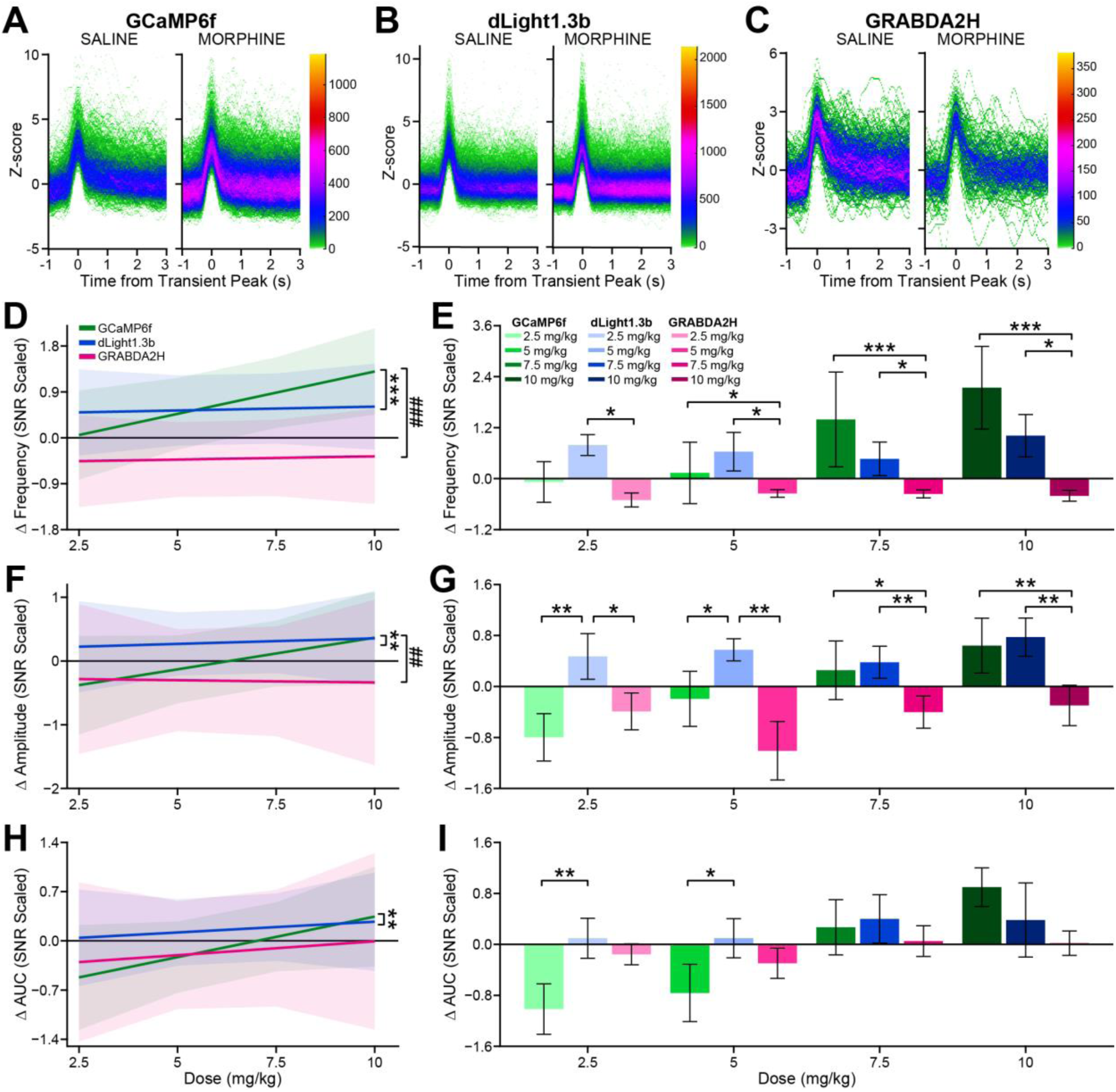
Comparison of spontaneous transient frequency, amplitude, and AUC between sensors. Color grade plots of overlaid individual transient traces from 10.0 mg/kg dose saline control and morphine treatment sessions for **A)** VTA DA GCaMP6f, **B)** NAcLS dLight1.3b, and **C)** NAcLS GRABDA2h. Color grade scale reflects the number of events overlapping at any given point. For between sensor comparisons, transient parameters were scaled to the mean and standard deviation of saline control sessions within each sensor to account for absolute differences between sensors and facilitate between sensor comparisons. **D)** Dose response slopes for the change in transient frequency with morphine administration relative to matched saline control sessions by sensor. **E)** Mean change in transient frequency by sensor across morphine doses. **F)** Dose response slopes for the change in transient amplitude by sensor. **G)** Mean change in transient amplitude by sensor across morphine doses. **H)** Dose response slopes for the change in transient AUC by sensor. **I)** Mean change in transient AUC by sensor across morphine doses. * <0.05; **,^##^<0.01; ***,^###^<0.001

#### 3.5.1. Transient Frequency

To compare effects of morphine on normalized transient frequency between sensors, we fit a linear mixed effects model to estimate the change in transient frequency during each morphine session from its matched saline session to examine the effects of sensor, dose, and time bin. Although there was a main effect of time bin (Χ^2^ = 42,12, *p =* 0.017), there were no interactions between dose and time bin (Χ^2^ = 12.61, *p =* 0.858), sensor and time bin (Χ^2^ = 32.73, *p =* 0.711), or dose, time bin, and sensor (Χ^2^ = 22.28, *p =* 0.980). Therefore, average change in transient frequency due to morphine treatment was averaged across the whole session. Overall, change in transient frequency depended on dose (Χ^2^ = 47.58, *p* < 0.0001), sensor (Χ^2^ = 8.15, *p =* 0.017), as well as an interaction between dose and sensor (Χ^2^ = 57.13, *p* < 0.0001).

To evaluate trends in dose-response effects, we fit simple slopes to quantify the change in transient frequency across doses for each sensor (Fig. 5D). GCaMP6f showed a robust positive relationship between morphine dose and transient frequency (Slope = 0.166, SE = 0.016). However, there were not dose-dependent effects on frequency with dLight1.3b (Slope = 0.015, SE = 0.016) or GRABDA2h (Slope = 0.013, SE = 0.018), and the slopes of these sensors were significantly lower than GCaMP6f (dLight1.3b: t(1876) = 6.66, *p* < 0.0001; GRABDA2h: t(1876) = 6.37, *p* < 0.0001), indicating reduced differentiation of morphine dose by transient frequency.

The direction of change in transient frequency differed across sensors. Following morphine treatment, transient frequency increased with GCaMP6f and dLight1.3b relative to saline sessions, but decreased GRABDA2h transient frequency (Fig. 5E). GCaMP6f showed more frequent transients in response to morphine compared to GRABDA2h at every dose higher than 2.5 mg/kg (5 mg/kg: t(1876) = 2.37, *p =* 0.018, d = 0.76; 7.5 mg/kg: t(1876) = 3.37, *p =* 0.0008, d = 1.09; 10 mg/kg: t(1876) = 4.28, *p* < 0.0001, d = 1.41). Similarly, change in transient frequency was significantly greater with dLight1.3b compared to GRABDA2h at all morphine doses (2.5 mg/kg: t(1876) = 2.47, *p =* 0.014, d = 0.81; 5 mg/kg: t(1876) = 2.54, *p =* 0.011, d = 0.82; 7.5 mg/kg: t(1876) = 2.55, *p =* 0.011, d = 0.82; 10 mg/kg: t(1876) = 2.50, *p =* 0.012, d = 0.83). Across all four doses, no significant differences between GCaMP6f and dLight1.3b were observed (all p > 0.06).

#### 3.5.2. Transient Amplitude

The change in transient amplitude with morphine treatment also revealed differences in sensor performance. Change in normalized transient amplitude depended on morphine dose (Χ^2^ = 17.89, *p* < 0.0001) and sensor (Χ^2^ = 10.34, *p =* 0.006), as well as an interaction between dose and sensor (Χ^2^ = 14.32, *p =* 0.001). Because there were no main effects of time bin (Χ^2^ = 21.85, *p =* 0.292) or interactions between dose and time bin (Χ^2^ = 10.11, *p =* 0.950), sensor and time bin (Χ^2^ = 42.17, *p =* 0.295), or dose, time bin, and sensor (Χ^2^ = 27.90, *p =* 0.885), change in transient amplitude for each morphine session from its matched saline session was averaged across the whole session.

The relationships between morphine dose and change in transient amplitude were qualitatively similar to the changes in frequency. GCaMP6f again showed a positive relationship between dose and amplitude (Slope = 0.099, SE = 0.018), while the relationship did not depend on dose for dLight1.3b (Slope = 0.018, SE = 0.017) or GRABDA2h (Slope = -0.007, SE = 0.032; Fig. 5F). The slope observed with GCaMP6f was significantly greater compared to both dLight1.3b (t(1462) = 3.36, *p =* 0.002) and GRABDA2h (t(1462) = 2.95, *p =* 0.009), while slopes did not differ between dLight1.3b and GRABDA2h (t(1462) = 0.69, *p =* 0.770).

The magnitude of changes in transient amplitude differed across the three sensors (Fig. 5G), but here differences between GCaMP6f and dLight1.3b were also evident. GCaMP6f amplitude decreased with the lowest dose of morphine, then increased in a dose-dependent manner. dLight1.3b displayed consistent elevations in transient amplitude across all four doses, while GRABDA2h showed consistent decreases. At the lower two doses of morphine, there was a dissociation between GCaMP6f and dLight1.3b, with GCaMP6f transient amplitude decreasing and dLight1.3b increasing (2.5 mg/kg: t(1462) = -3.09, *p =* 0.002, d = -0.49; 5 mg/kg: t(1462) = -2.28, *p =* 0.023, d = - 0.32). Both sensors showed increased amplitude that did not differ from each other at higher doses (7.5 mg/kg: t(1462) = -1.11, *p =* 0.266, d = -0.16; 10 mg/kg: t(1462) = 0.06, *p =* 0.956, d = 0.01). At higher, but not lower, doses, GCaMP6f showed larger amplitude transients than GRABDA2h (7.5 mg/kg: t(1462) = 2.12, *p =* 0.034, d = 0.346; 10 mg/kg: t(1462) = 2.85, *p =* 0.004, d = 0.57). Like frequency, the change in transient amplitude at dLight1.3b sensors was significantly higher than GRABDA2h across all doses (2.5 mg/kg: t(1462) = 2.20, *p =* 0.028, d = 0.41; 5 mg/kg: t(1462) = 2.85, *p =* 0.004, d = 0.46; 7.5 mg/kg: t(1462) = 3.07, *p =* 0.002, d = 0.51; 10 mg/kg: t(1462) = 2.81, *p =* 0.005, d = 0.56).

#### 3.5.3. Transient Area Under the Curve

To capture differences in morphine effects on overall shape of transients across sensors, we also analyzed normalized change in transient AUC. We observed a significant main effect of dose (Χ^2^ = 36.12, p < 0.0001) and an interaction between dose and sensor (Χ^2^ = 13.56, *p =* 0.001). No other significant main effects or interactions were observed (all *p* > 0.14).

All three sensors showed a positive relationship between morphine dose and change in transient AUC (Fig. 5H). GCaMP6f showed a stronger relationship between dose and AUC (Slope = 0.115, SE = 0.017) than dLight1.3b (Slope = 0.030, SE = 0.016; t(1462) = 3.59, *p =* 0.001) but was not different from GRABDA2h (Slope = 0.039, SE = 0.030; t(1462) = 2.18, *p =* 0.076). No differences in slopes were observed between GRABDA2h and dLight1.3b (t(1462) = -0.253, *p =* 0.965).

The magnitude of changes in transient AUC across sensors was most apparent at lower doses of morphine (Fig. 5I). Transient AUC was suppressed with GCaMP6f compared to dLight1.3b at morphine doses of 2.5 mg/kg (t(1462) = -3.18, *p =* 0.001, d = -0.47) and 5 mg/kg (t(1462) = -2.25, *p =* 0.024, d = -0.29). No differences between GCaMP6f and dLight1.3b were observed with 7.5 and 10 mg/kg.

#### 3.5.4. Summary of Morphine Effects on Transients Across Sensors

Only VTA_DA_ GCaMP6f showed a positive linear relationship between morphine dose and transient frequency as well as amplitude. NAcLS dLight1.3b responded with increases in transient frequency and amplitude, but in a relatively dose-independent manner. NAcLS GRABDA2h responses not only did not depend on morphine dose but were characterized by consistent decreases in signal. As a substantial literature confirms that morphine increases the release of dopamine in NAc, the reduction in transients measured by the GRABDA2h sensor is unexpected. This sensor differs from dLight1.3b in both its affinity for dopamine as well as the on- and off-kinetics of its binding (Table 1). To elucidate if morphine-driven effects on dopamine transients were impacted by sensor differences in affinity and kinetics, we examined changes in the mean and variation of whole data streams, as these sensor characteristics might reveal more generalized signal changes than reflected in transient events.

**Table 1.**
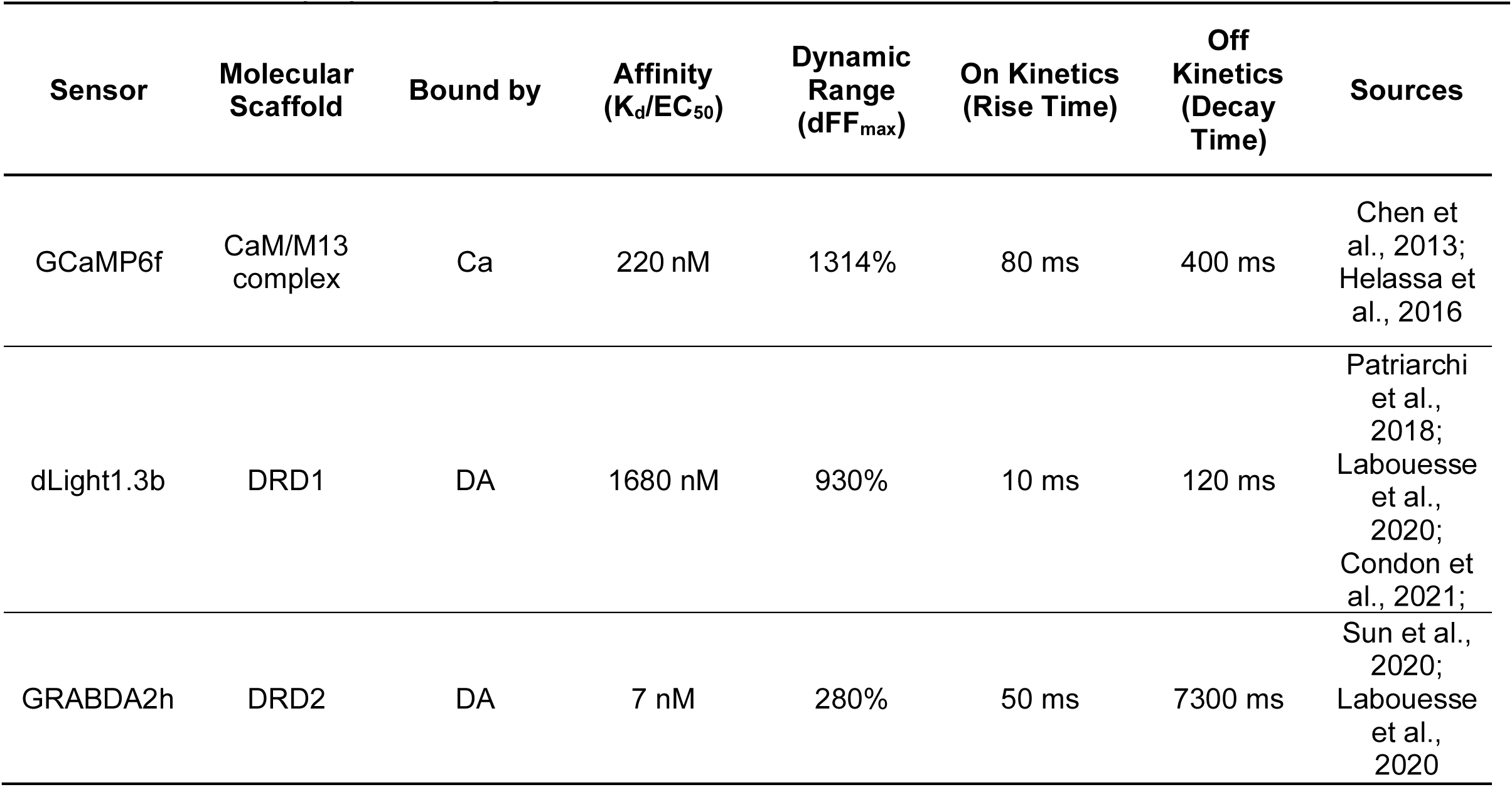
Sensor affinity, dynamic range, and kinetics.

| Sensor | Molecular Scaffold | Bound by | Affinity (K<sub>d</sub>/EC<sub>50</sub>) | Dynamic Range (dFF<sub>max</sub>) | On Kinetics (Rise Time) | Off Kinetics (Decay Time) | Sources |
| --- | --- | --- | --- | --- | --- | --- | --- |
| GCaMP6f | CaM/M13 complex | Ca | 220 nM | 1314% | 80 ms | 400 ms | Chen et al., 2013; Helassa et al., 2016 |
| dLight1.3b | DRD1 | DA | 1680 nM | 930% | 10 ms | 120 ms | Patriarchi et al., 2018; Labouesse et al., 2020; Condon et al., 2021; |
| GRABDA2h | DRD2 | DA | 7 nM | 280% | 50 ms | 7300 ms | Sun et al., 2020; Labouesse et al., 2020 |

### 3.6. Whole Session Quantification

We compared the mean and mean absolute deviation (MAD) of unfiltered subtracted data streams – independent of transients – between morphine treatment sessions and their matched saline control sessions separately in animals expressing GCaMP6f, dLight1.3b, and GRABDA2h sensors. Mean traces and confidence intervals across all subjects by sensor and injection type (saline vs morphine) for the highest dose of 10.0 mg/kg are displayed in Figure 6 A-C.

**Figure 6.**
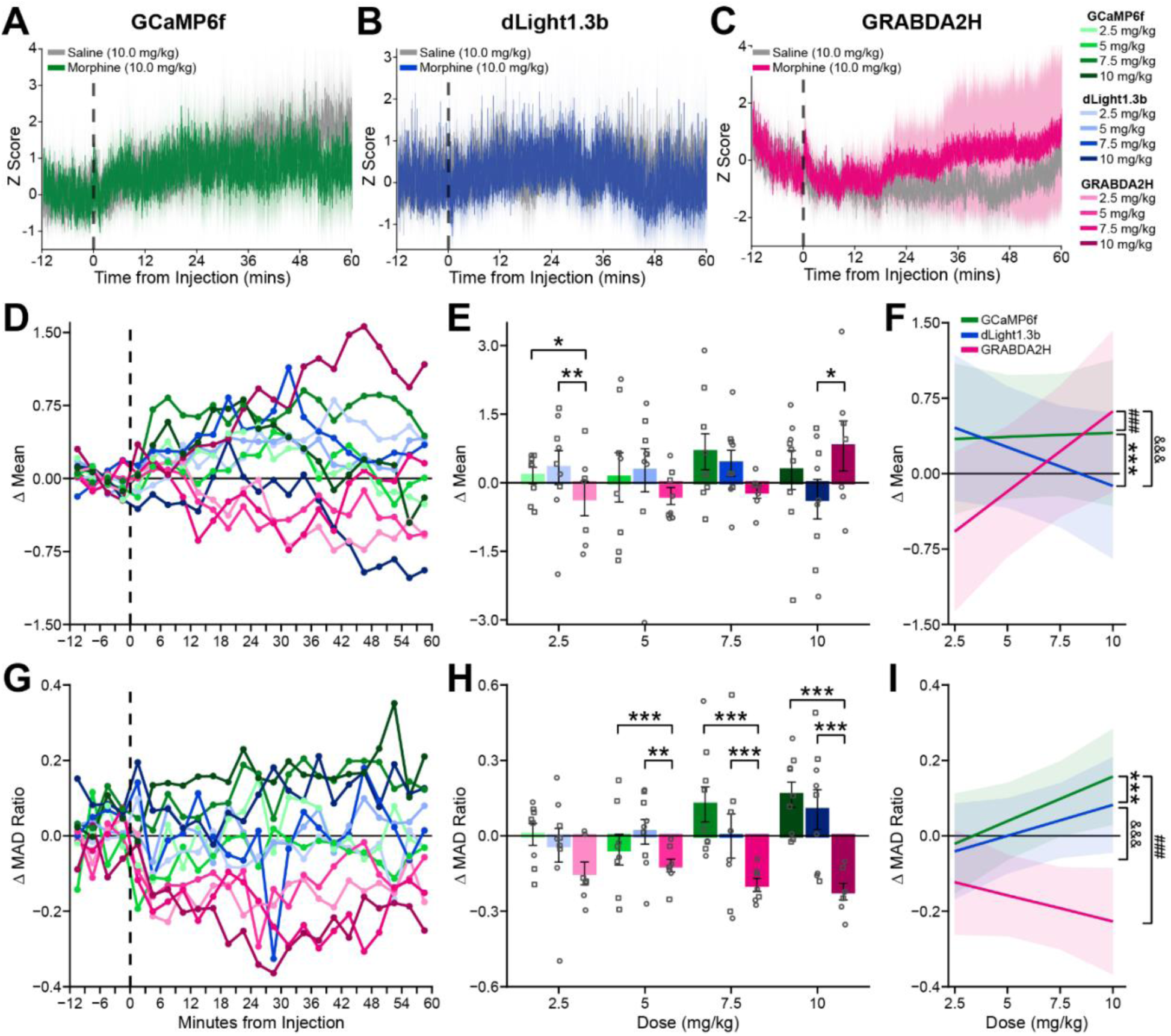
Comparison of whole stream mean and mean absolute deviation (MAD) between sensors. Average whole session traces with shaded confidence intervals from 10.0 mg/kg dose saline control and morphine treatment sessions for **A)** VTA DA GCaMP6f, **B)** NAcLS dLight1.3b, and **C)** NAcLS GRABDA2h. For between sensor whole stream comparisons of mean and MAD, data were analyzed in 3-minute time bins across the session. **D)** Change in stream mean by time bin, sensor, and morphine dose relative to matched saline control sessions. **E)** Mean change in whole stream mean by sensor and morphine dose. **F)** Dose response slopes for change in whole stream mean by sensor. **G)** Change in stream MAD by time bin, sensor, and morphine dose relative to matched saline control sessions. **H)** Mean change in whole stream MAD by sensor and dose. **I)** Dose response slopes for change in whole stream MAD by sensor. * <0.05; **<0.01; ***, ^&&&^, ^###^ <0.001

#### 3.6.1. Whole Session Signal Mean Comparison

For each session, the mean overall signal was determined in three-minute time bins. Change values were calculated to compare each treatment session to its matched saline control bin-by-bin (Fig. 6D). Overall, GCaMP6f and dLight1.3b produced comparable results, while effects with GRABDA2h diverged in a dose-dependent manner. We observed significant interactions between dose and sensor (Χ^2^ = 139.80, *p* < 0.0001) and between dose, sensor, and time bin (Χ^2^ = 66.40, *p =* 0.003). Pairwise comparisons revealed that with 2.5 mg/kg, the change in signal mean was significantly lower with GRABDA2h compared to both GCaMP6f (t(1855) = 2.34, *p =* 0.017) and dLight1.3b (t(1855) = 2.68, *p =* 0.008; Fig. 6E). By contrast, with the highest dose of 10.0 mg/kg, the mean signal with GRABDA2h trended higher compared to dLight1.3b (t(1855) = -1.92, *p =* 0.055). No differences were observed between GCaMP6f and dLight1.3b across all doses (all *p* > 0.15).

Comparison of relationships between morphine dose and mean signal showed different patterns between GRABDA2h and the other sensors (Fig. 6F). While both GCaMP6f and dLight1.3b showed minimal effects of dose on mean signal (GCaMP6f: Slope = 0.008, SE = 0.014; dLight1.3b: Slope = -0.077, SE = 0.013), GRABDA2h showed a positive relationship (Slope = 0.16, SE = 0.015). The slope of GRABDA2h was significantly greater than both GCaMP6f (t(1855) = -7.43, *p* < 0.0001) and dLight1.3b (t(1855) = -11.79, *p* < 0.0001). The slope of dLight1.3b was more negative than GCaMP6f (t(1855) = 4.53, *p =* < 0.0001).

#### 3.6.2. Whole Session Signal Mean Absolute Deviation (MAD)

To quantify changes in the variation of the whole session, the mean absolute deviation (MAD) was calculated in three-minute time bins. MAD values for morphine sessions were scaled (MAD Ratio) to their matched saline sessions within sensor to facilitate cross-sensor comparison. Generally, morphine treatment increased the MAD ratio with GCaMP6f and dLight1.3b but caused reductions with GRABDA2h (Fig. 6G). We observed significant main effects of dose (Χ^2^ = 46.24, p < 0.0001), sensor (Χ^2^ = 20.57, *p* < 0.0001), as well as significant interactions between dose and sensor (Χ^2^ = 96.08, *p* < 0.0001), and sensor and time bin (Χ^2^ = 54.39, *p =* 0.042). Pairwise comparisons revealed that GRABDA2h exhibited reductions in MAD ratio that were, with the exception of the 2.5 mg/kg dose of morphine, markedly different from GCaMP6f and dLight1.3b (Fig. 6H). Specifically, change in MAD ratio with GRABDA2h was significantly lower than GCaMP6f (5.0 mg/kg: t(1855) = 3.73, *p =* 0.0002, d = 1.03; 7.5 mg/kg: t(1855) = 5.50, *p* < 0.0001, d = 1.52; 10.0 mg/kg: t(1855) = 7.03, *p* < 0.0001, d = 2.01) and dLight1.3b (5.0 mg/kg: t(1855) = 3.00, *p =* 0.003, d = 0.83); 7.5 mg/kg: t(1855) = 4.42, *p* < 0.0001, d = 1.22; 10.0 mg/kg: t(1855) = 5.66, *p* < 0.0001, d = 1.62). No differences in MAD ratio changes were observed across doses between GCaMP6f and dLight1.3b (all *p* > 0.15).

Like mean signal, the comparison of relationships between morphine dose and change in MAD ratio showed very different patterns between GRABDA2h and the other sensors (Fig. 6I). Both GCaMP6f and dLight1.3b sensors showed positive relationships (GCaMP6f: Slope = 0.024, SE = 0.003; dLight1.3b: Slope = 0.016, SE = 0.003) that did not differ (t(1855) = 1.98, *p =* 0.117). In contrast, the relationship between dose and change in MAD ratio was negative with GRABDA2h (Slope = -0.014, SE = 0.003), significantly differing from both GCaMP6f (t(1855) = 9.38, *p* < 0.0001) and dLight1.3b (t(1855) = 7.69, *p* < 0.0001). Combined, these results show that, when using GRABDA2h, there was an overall increase in whole signal data stream combined with reduced variability after morphine injection.

## 4. DISCUSSION

Acute, systemic opioid administration causes an increase in the firing rate and burst firing of presumed dopamine neurons (Jalabert et al., 2011). It also causes increased dopamine release in the NAc as measured with microdialysis (Di Chiara & Imperato, 1988; Pothos et al., 1991; Rougé-Pont et al., 2002) and fast-scan cyclic voltammetry (Fox et al., 2016; Vander Weele et al., 2014). With increasing adoption of in vivo fiber photometry to elucidate neural substrates of behavior, it is essential to characterize fluorescent sensor performance across a range of experimental conditions and pharmacological manipulations. Yet much of the validation of sensors has been conducted in vitro or in response to salient, non-pharmacological stimuli ((Patriarchi et al., 2018; Sun et al., 2020) but see also (Salinas et al., 2023)). Acute, systemic morphine is an ideal pharmacological manipulation to compare sensor performance given the well-established response of the mesolimbic dopamine system. Responses to escalating doses of morphine across VTA_DA_ GCaMP6f, NAcLS dLight1.3b, and NAcLS GRABDA2h sensors allowed for assessment of sensor performance compared with prior research findings. We characterized shifts in DA to capture fast (i.e. transients) and slow (i.e. whole session mean and MAD) dynamics and made direct comparisons across sensors. While all three sensors detected spontaneous transient events during saline-control sessions and pre-morphine injection baseline periods, the performance of GRABDA2h following morphine injection substantially diverged from both GCaMP6f and dLight1.3b.

Exogenous opioids are positively reinforcing through the disinhibition of VTA dopamine neurons. Systemic morphine treatment increases the firing rate of presumed dopamine neurons, modulating distinct aspects of activity, including increasing burst rate and the number of spikes in a burst (Chen et al., 2015; Georges et al., 2006; Grant & Sonti, 1994; Jalabert et al., 2011; Johnson & North, 1992; Matthews & German, 1984; Nowycky et al., 1978; Wu et al., 2025). Further, the elimination of mu-opioid receptors in the VTA abolishes opioid reinforcement and opioid-induced DA release in the NAc averaged over many minutes (Chaudun et al., 2024). Here, consistent with the literature, we found dose-dependent increases in the frequency, amplitude, and AUC of GCaMP6f transients in VTA. While all doses increased GCaMP6f transient frequency, morphine exhibited a bidirectional effect on amplitude and AUC; lower doses decreased amplitude and AUC but higher doses increased amplitude and AUC relative to saline. How changes in the parameters of calcium dynamics (e.g. frequency versus amplitude) map on to electrophysiological (e.g. burst rate versus number of spikes in a burst) signatures is critical. While some studies average responses over minutes (as in Chaudun et al. 2024, for example), here, effects were explored on the foundation of identifying transient events that last ∼1-2 seconds. Different time courses for analyses may contribute to diverging results between electrophysiological recordings and GCaMP-based fiber photometry.

The dorsolateral VTA_DA_ neurons targeted here project to the NAcLS (Breton et al., 2019). While some work has identified high fidelity between DA cell body activity and release in the NAc (Lee et al., 2020), recent work has also suggested that processes at dopamine terminals may modulate DA release dynamics beyond cell body activity (Touponse et al., 2026). Using the D1-based sensor dLight1.3b, we found that systemic morphine increased frequency and amplitude of DA transients. Results were largely similar to those observed with GCaMP6f transients in the VTA, however dLight1.3b transients did not show dose-dependency. Morphine can alter aspects of cholinergic neuron function in the NAc (Zucca et al., 2025) which could, in turn, represent a mechanism by which measures of DA release are tuned by terminal processes. Given the heterogeneity of dopamine neurons, their responses to opioids (Morison et al., 2025), and their projections to distinct NAc subregions (Miranda-Barrientos et al., 2021), it will be critical to continue to characterize optical sensors across NAc and striatal subregions (as in Salinas et al. (2023) for responses to cocaine, example) for their response to systemic treatment.

We also used GRABDA2h to measure opioid-induced transients in the NAcLS. GRABDA2h captures changes evoked by both rewarding and aversive physiological and behavioral stimuli (Sun et al., 2020) that are consistent with the literature. Here, though, unlike the results with dLight1.3b, we found decreases in frequency, amplitude, and AUC of dopamine transients relative to saline control. The GRABDA2h sensor is scaffolded on the high affinity D2 receptor and is known to have high sensitivity but slow on/off kinetics. As such, it is possible that the increase in dopamine activity induced by morphine – as observed in our experiments using GCaMP6f and dLight1.3b and consistent with electrophysiological and voltammetry recordings – overwhelmed the ability of the GRABDA2h sensor to maintain fidelity to events on a sub-second timescale. Suppression of phasic dopamine release by exogenous opioid (e.g. fentanyl; (Culver et al., 2026)) observed using the D2-based sensor GRABDA1m was recently reported. Similar to our results where we observed a slow increase in fluorescence over minutes from morphine injection, Culver et al. also reported an increase in ‘tonic’ dopamine after fentanyl administration. Distinguishing between saturation/floor/ceiling effects and decreases in transient dynamics will be a continued challenge as sensors are refined (Roshgadol et al., 2025; Zhuo et al., 2024).

Analysis methods selected to normalize data streams for the detection and quantification of transient events also merit consideration. While normalization techniques such as Z-scoring can be critical to standardize the scale of recordings between subjects, the reference period used for normalization must be carefully chosen. If the mean and standard deviation used for normalization include data obtained during pharmacological treatment, shifts in signal range and variation may be obscured or misrepresented if they contribute to the normalization itself (Wallace et al., 2025). Additionally, transient event detection methods can be skewed by both local and global fluctuations in the data stream. Methods that utilize sliding windows or prominence-based thresholds may therefore over or under detect events when applied to analyze the effects of pharmacological treatments. Additionally, these methods limit comparisons across multiple recording sessions as event inclusion thresholds based on sliding windows or rolling prominence estimates are inconsistently applied both within and between recording sessions. The default values of our analysis pipeline have been selected to maintain threshold consistency across experimental conditions (Donka et al., 2025).

Compared with GCaMP6f and dLight1.3b, spontaneous transient signals with GRABDA2h measured prior to any drug treatment were less frequent, of lower amplitude, with significantly slower off-kinetics. It is possible that these intrinsic properties of the sensor compromised its representation of underlying signal dynamics with fidelity, in effect smoothing the phasic nature of DA transients over time. By examining unfiltered signals across entire recording sessions, we observed another unique property of GRABDA2h relative to the other sensors – a slow signal increase. Apparent sustained increases in signal may result from averaging individual phasic release events across increasingly longer time-bins (Owesson-White et al., 2012). However, we also observed a significant reduction in the variability of unfiltered signal, quantified as MAD ratio, after morphine administration, aligning with the observed reduction in frequency and amplitude of transient events. Given that GRABD2h has higher affinity for dopamine and slower kinetics than dLight1.3b, the increase in overall signal coupled with the decrease in MAD ratio is consistent with an interpretation that the GRABDA2h signal after morphine treatment reflects saturation of the sensor. In contrast, both GCaMP6f and dLight1.3b showed increased frequency and amplitude of transient events, reflected in the overall increase in MAD ratio, again opposite to the patterns observed with GRABDA2h. Additional studies will need to be conducted to determine how pharmacological treatments modulate responses to natural stimuli to determine whether transient events themselves as well as the ability of transient events to synchronize to behavioral events are altered.

These results suggest that the use of biosensors to assay DA release and underlying dynamics should be done with caution, with the inclusion of positive and negative controls for pharmacological responses, ensuring that sensor selection is suited for the region of study and experimental paradigm. Consideration of sensor dynamics (sensitivity, dynamic range, kinetics) and time scale of the phenomenon under question is critical. Use of a sensor that fails to capture dynamics may result in not just null findings but illusory conclusions that do not reflect the actual activity and state of the underlying system. Additionally, the inclusion of positive and negative controls are critical for sensor validation in the experimental context, limiting the risk of data interpretation misled by sensor performance. While some sensors may be sufficient for capturing dynamics evoked by natural stimuli or transient artificial stimuli such as electrical stimulation, they may falter under the more extreme and long-lasting effects of pharmacological manipulations. Ongoing studies to characterize the basic properties of sensor behavior in vivo under multiple pharmacological and behavioral paradigms is critical to establish consistent biological interpretations of shifts in fluorescence as neural activity.

## Data Availability Statement

Processed data and analysis tables are stored at https://zenodo.org/uploads/20752722 (DOI: 10.5281/zenodo.20752722). Analysis scripts are available at https://github.com/rdonka/SensorSensibility. Raw data files are available upon request.

## Author Contributions

**RMD**: Conceptualization, Data curation, Methodology, Software, Formal analysis, Visualization, Writing – original draft. **MKL**: Data curation, Software, Writing – review and editing. **MFR**: Conceptualization, Supervision, Funding acquisition, Writing – review and editing. **JDR**: Conceptualization, Project administration, Funding acquisition, Writing – review and editing.

